# Structured cross-omics interaction discovery with a triple-graph model

**DOI:** 10.64898/2026.09.10.749540

**Authors:** Jiaao Yu, Huang Lin, Shuo Chen

**Author notes:** **Corresponding author** Correspondence to Huang Lin and Shuo Chen. Contributing authors. These authors contributed equally to this work (Jiaao Yu, Huang Lin).

## Abstract

Multi-omics analyses often yield fragmented pairwise associations that obscure coordinated relationships among molecular features. We developed TriGer, a triple-graph framework that identifies many-to-many cross-omics modules by combining cross-layer associations with dependency structures within each layer. In simulations with sparse or nested signals, TriGer recovered planted modules while balancing sensitivity and specificity. In inflammatory bowel disease, it identified subtype-associated metabolite–transcript modules; in colorectal cancer, it identified genus–metabolite modules whose organization was attenuated in cancer. TriGer provides an interpretable approach for studying coordinated cross-omics structure in high-dimensional molecular data.

## Background

High-throughput molecular profiling technologies now enable biological systems to be characterized across multiple molecular layers, including the genome, epigenome, transcriptome, proteome, metabolome, and microbiome [1]. These advances have shifted biomedical research from single-layer analyses towards integrative, systems-level investigation [2, 3]. Molecular layers do not operate independently; rather, they are linked through regulatory, biochemical and ecological interactions that together shape cellular function and organismal phenotypes [4]. Genetic variation can propagate through transcriptional, metabolic, and microbial processes, and the resulting phenotypes often reflect coordinated perturbations across these layers rather than changes in any one layer alone [5, 6]. Accordingly, multi-omic integration has become central to efforts to understand biological organization, disease heterogeneity and the mechanisms that connect molecular variation to clinical outcomes.

The need for integrative analysis is especially acute in complex diseases, in which pathological processes emerge from interactions among many biological components operating at different scales. Host–microbiome systems provide a particularly clear example. In these systems, microbial communities, host gene regulation, and metabolism are tightly coupled through microbial metabolites, immune signalling and host physiological responses [7]. In inflammatory bowel disease (IBD), for example, disease activity has been associated with coordinated alterations in the gut microbiome, host transcriptional programs, and metabolic pathways, underscoring that disease-relevant biology is distributed across inter-connected molecular layers rather than localized to a single source [8]. Large longitudinal efforts such as the Integrative Human Microbiome Project (iHMP) have further reinforced this perspective by generating rich, multi-layer datasets paired with clinical and environmental measurements [8, 9]. These resources create unprecedented opportunities to identify cross-omics patterns that explain disease progression, inter-individual variation and system-level organization.

Realizing that potential, however, remains difficult. Multi-omic datasets are typically high dimensional, with the number of measured features far exceeding the number of samples, making inference statistically unstable and computationally demanding [10]. Signals of interest are often weak, distributed across many variables and obscured by measurement noise, batch effects and substantial between-subject heterogeneity [11, 12]. As a result, feature-by-feature analyses face two related limitations. First, they are frequently underpowered, and stringent multiple-testing correction often leaves only the strongest marginal associations. Second, even when numerous pairwise associations are detected, the resulting signal can be diffuse and difficult to interpret, appearing as scattered patterns across a high-dimensional association map rather than as coherent biological organization. More fundamentally, biological effects in complex disease are rarely isolated to single variables; instead, they are expressed through coordinated groups of features acting within and across molecular layers. Distinguishing such structured biological signal from random or fragmented correlation therefore remains a major challenge in multi-omics analysis [13].

Existing analytical approaches address only part of this challenge because they tend to emphasize either global shared variation or isolated cross-omics associations, but not the structured many-to-many relationships through which biological processes are often organized. Dimension-reduction or matrix-factorization methods, including principal-component-based approaches [14, 15] and multi-omics factor models such as MOFA [16], FABIA [17], and JIVE[18], are primarily driven by dominant sources of variation. Because these methods operate directly on high-dimensional data matrices, they are susceptible to noise amplification and, when the number of molecular features far exceeds the sample size, often struggle to identify stable shared components because the optimization problem becomes ill-conditioned. Just as importantly, they do not explicitly target the cross-omics correlation structure itself, so coordinated cross-layer signals can be absorbed into modality-specific components and remain undetected. Clustering-based integration approaches, including sparse canonical correlation analysis [19, 20] and DIABLO [21], move closer to the goal of identifying relationships between omics layers, but they still typically focus on pairwise or one-to-one patterns and rely on continuous weight vectors rather than discrete module structures. As a result, within-layer organization is only weakly represented, variables from distinct biological modules can be mixed together, and false discoveries become more likely when the true signal is sparse. More broadly, most current approaches still frame cross-omics discovery as a feature-by-feature problem, asking whether one variable in one layer is associated with one variable in another. That formulation can be statistically inefficient when the underlying biology is many-to-many: weak but coordinated signals spread across groups of features may fail to reach significance individually even when, taken together, they define a reproducible cross-omics program.

Taken together, these limitations point to a clear methodological gap. What is needed is an integrative framework that does not force a choice between cross-omics association and within-omics structure, but instead models them jointly so that groups of features are identified because they form coherent higher-order patterns across molecular layers. Such a framework should aggregate weak, distributed signals into stable modules, preserve the internal organization of each data type, and provide outputs that remain biologically interpretable. By shifting the inferential target from isolated pairs to interacting feature sets, it can borrow strength across aligned signals, reduce fragmentation caused by massive multiple testing, and improve sensitivity to structured but modest effects. Without this joint treatment of within-layer and cross-layer organization, important disease-relevant patterns are likely to remain fragmented across separate analyses or diluted by dominant but biologically less informative sources of variation.

To address this gap, we developed **Tri**ple-**G**raph Multi-omics Int**er**action Model (TriGer), an integrative framework that treats the primary unit of signal as a structured cross-omics module rather than an isolated association. TriGer starts from the premise that biologically meaningful interactions are often expressed as coordinated many-to-many relationships, in which groups of features in one molecular layer align with groups in another while also retaining coherent organization within each layer. It therefore combines evidence from cross-layer association and within-layer dependency in a single graph-based representation, allowing weak but concerted signals to accumulate at the module level rather than being evaluated only as disconnected pairs. This design yields outputs that are not only more stable in high-dimensional settings but also more directly interpretable biologically, because the recovered modules preserve both cross-omics coupling and the internal structure of each modality. In this way, TriGer provides a scalable framework for characterizing system-level molecular organization and for linking coordinated multi-omic programs to complex disease phenotypes.

## Results

### Overview of the analytical framework

TriGer identifies structured cross-omics modules in two stages. It first searches the cross-omics association matrix for dense blocks and then refines the selected features using within-omics correlation structure. The final output is therefore a pair of feature sets, one from each omics layer, together with the cross-omics block linking them.

This two-stage design preserves both cross-layer coupling and within-layer coherence, yielding modules that are more compact and interpretable than diffuse collections of pairwise associations. Full mathematical formulation, optimization details, theoretical support, and tuning-parameter selection are provided in the Methods section.

### Simulation benchmarking supports joint recovery of cross- and within-omics structure

Simulation analyses showed that TriGer more reliably recovered the targeted block structure than the comparison methods, particularly when signal was sparse or nested. We considered two benchmark settings. Scenario 1 contained one large planted cross-omics block together with a smaller nested block embedded within it. Scenario 2 was more challenging and contained three planted blocks with hierarchical overlap, including progressively smaller nested structures. In Table 1, these planted signals are labeled as the primary block and nested block(s), and performance is summarized using true positive rate (TPR) and true negative rate (TNR) for recovery of each block. Full simulation settings, including block sizes, correlation parameters, and data-generation details, are provided in the Methods section.

**Table 1:**
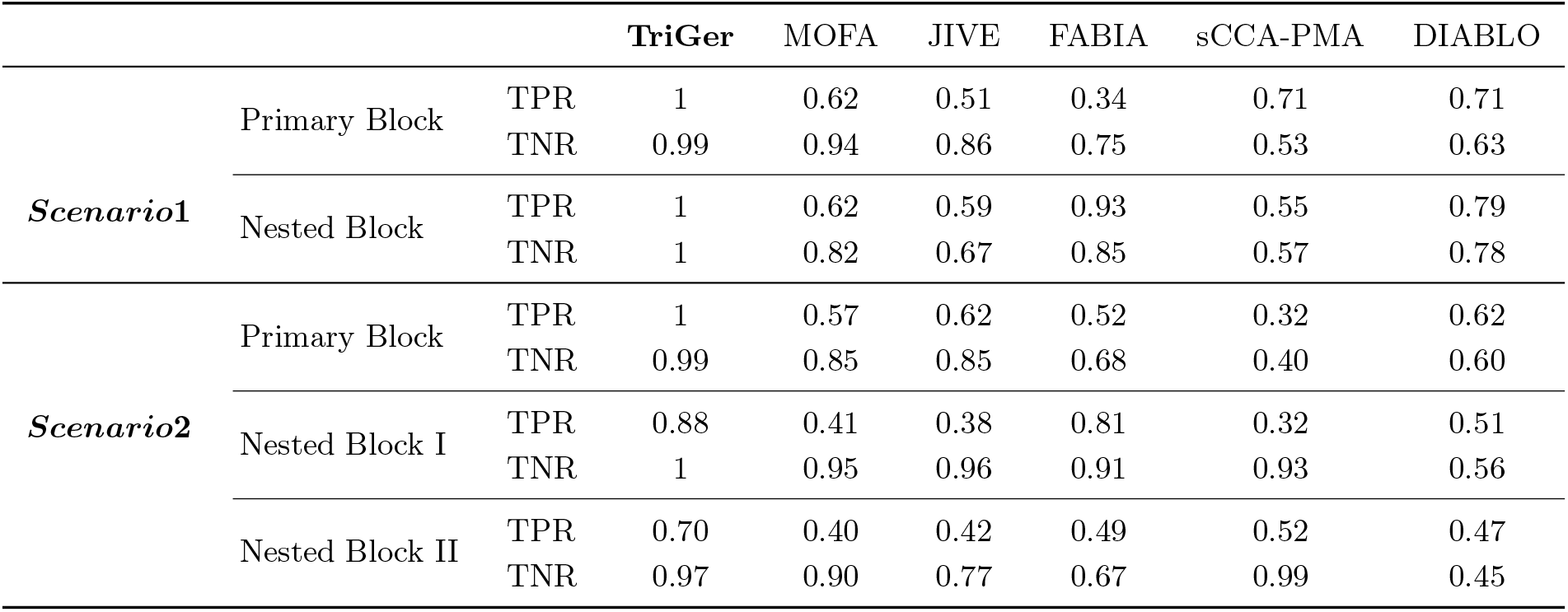
Recovery of planted cross-omics blocks in simulation benchmarks. Scenario 1 contains one primary block and one nested block, whereas Scenario 2 contains one primary block and two progressively smaller nested blocks. For each planted block, true positive rate (TPR) and true negative rate (TNR) summarize the sensitivity and specificity with which each method recovered the corresponding signal region.

Table 1 shows that TriGer was the only method to maintain both high sensitivity and high specificity across the planted blocks in both simulation scenarios. In Scenario 1, TriGer recovered both the primary block and the nested block almost perfectly (TPR = 1.00 for both; TNR = 0.99 and 1.00, respectively), indicating that it captured the large signal region as well as the embedded substructure without appreciable contamination. A similar pattern held in Scenario 2, where performance remained strong for the primary block (TPR = 1.00, TNR = 0.99) and degraded only modestly for the more difficult nested blocks (nested block 1: TPR = 0.88, TNR = 1.00; nested block 2: TPR = 0.70, TNR = 0.97). The competing methods failed in more specific ways. MOFA and JIVE were moderately competitive on the largest planted blocks, but both lost sensitivity once the target structure became smaller or more deeply nested. In Scenario 2, for example, MOFA and JIVE achieved TPR values of only 0.41 and 0.38 for nested block 1, and 0.40 and 0.42 for nested block 2, despite retaining relatively high TNR. This pattern suggests that these methods often captured broad shared variation while missing weaker embedded blocks. FABIA showed the opposite tendency. It sometimes recovered larger portions of a true block, particularly for nested structures (for example, TPR = 0.93 for the Scenario 1 nested block and 0.81 for Scenario 2 nested block 1), but this came at the cost of reduced specificity, with TNR values falling to 0.85 for the Scenario 1 nested block and 0.67 for Scenario 2 nested block 2. Thus, FABIA tended to over-extend detected modules by including variables outside the planted block. sCCA-PMA was particularly prone to false positive inclusion in the larger blocks. For the primary block in Scenario 1, it reached TPR = 0.71 but with TNR only 0.53, and for the primary block in Scenario 2 the corresponding values were 0.32 and 0.40. Even when specificity improved for smaller planted blocks, sensitivity remained limited. This behavior is consistent with the canonical-correlation objective selecting mixtures of signal and non-signal variables when no explicit block-density constraint is imposed. DIABLO yielded intermediate performance but was less stable across settings. In Scenario 1 it recovered part of both the primary and nested block (TPR = 0.71 and 0.79), but with only moderate specificity (TNR = 0.63 and 0.78). In Scenario 2, performance deteriorated further in the more challenging settings, especially for nested block 2, where TPR fell to 0.47 and TNR to 0.45. Taken together, these results indicate that methods not designed to recover dense cross-omics blocks with aligned within-omics structure either miss nested signal or recover it at the expense of substantial false positive inclusion, whereas TriGer preserves a more favorable balance between the two.

### Structured metabolite–transcript modules distinguish IBD subtypes

We next applied TriGer to the inflammatory bowel disease (IBD) cohort from the Inflammatory Bowel Disease Multiomics Database (IBDMDB) [8]. The analysis included 76 individuals with matched metabolomics and transcriptomics profiles: 32 with Crohn’s disease (CD), 22 with ulcerative colitis (UC), and 22 non-IBD controls. To retain signals that were strong in at least one clinical group, we combined subgroup-specific metabolite–transcript association evidence into a single screening matrix using the strongest signal observed across CD, UC, and control samples (see Methods). This strategy preserves subgroup-restricted associations without requiring concordance across all three groups. As shown in Fig. 1, the aggregated matrix appears largely sparse before reordering, with little immediately visible structure. After TriGer-guided reordering of rows and columns, however, a prominent dense block emerges in the top-left corner, corresponding to a compact set of metabolites and transcripts with consistently strong cross-omics associations in at least one subgroup. We then projected the selected features back onto each subgroup-specific matrix to determine whether the recovered structure was shared across groups or enriched in a particular disease subtype.

**Fig. 1:**
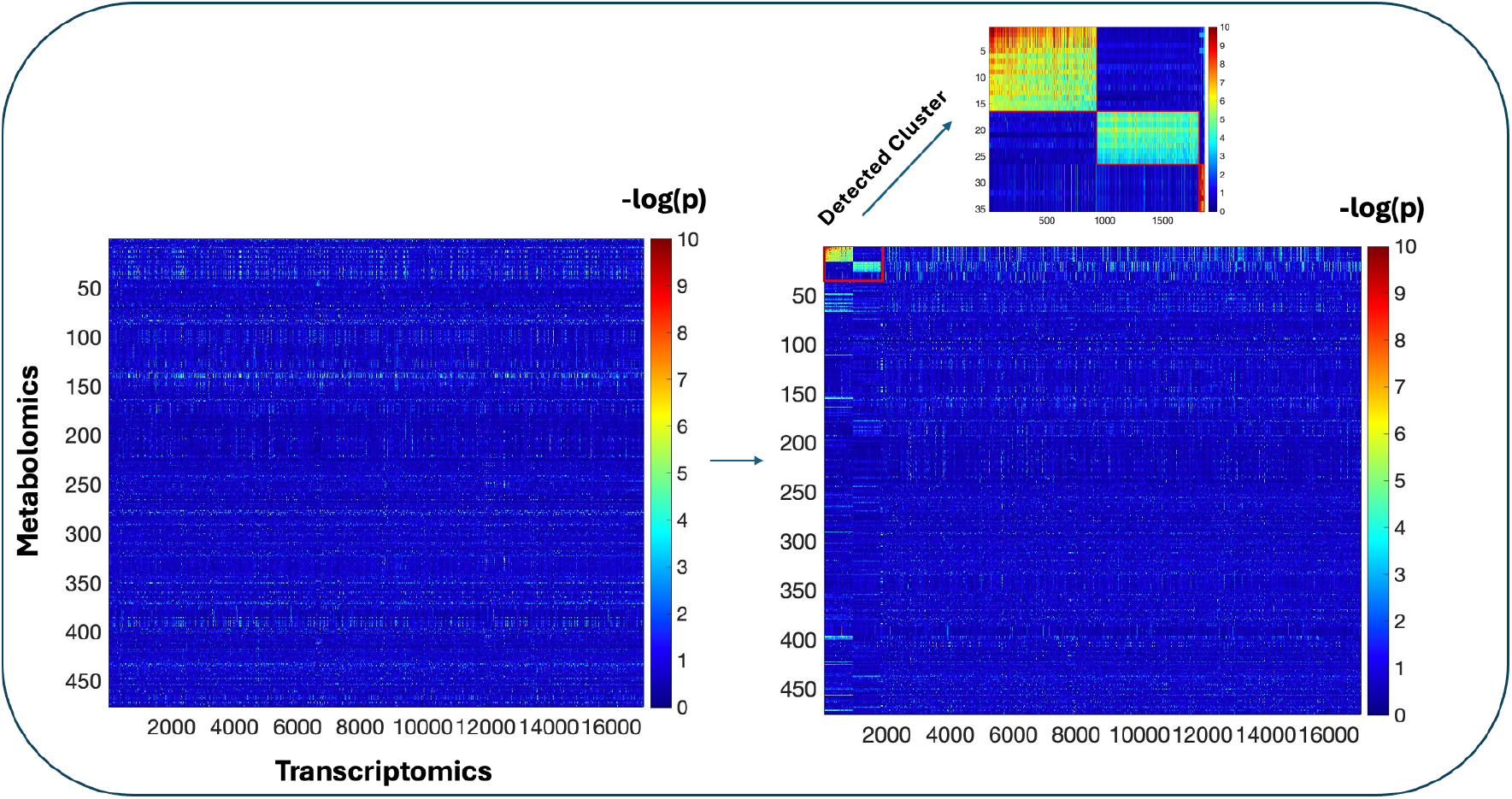
TriGer reveals a dense metabolite–transcript association block in IBD. Aggregated cross-omics association matrix constructed by computing group-specific correlation matrices for Crohn’s disease (CD), ulcerative colitis (UC), and non-IBD controls separately, applying a − log_10_(*p*) transformation to each, and retaining the maximum value across three groups for each feature pair. Each element represents the statistical significance of the association between a metabolite (rows, *n* = 476) and a transcript (columns, *n* = 17,026), with warmer colors indicating stronger associations. The left panel shows the original matrix, and the right panel shows the same matrix after TriGer-guided reordering of rows and columns, which exposes a prominent dense block in the top-left corner. The inset highlights the detected module, corresponding to a compact set of metabolites and transcripts with coordinated subgroup-specific associations.

To visualize the recovered structure within each disease group, we assembled a joint block matrix containing metabolite–metabolite correlations, transcript–transcript correlations, and metabolite–transcript associations restricted to the variables selected by TriGer. This representation makes it possible to inspect within-omics organization and cross-omics coupling in a common coordinate system. Fig. 2 shows that the recovered module is not expressed uniformly across the three groups. UC samples display the clearest block organization, with stronger and more spatially coherent metabolite–transcript coupling accompanied by pronounced within-omics structure (Modules 1 and 2). By contrast, the analogous regions in CD and non-IBD controls are weaker and less sharply organized. These differences indicate that the detected module captures cross-omics coordination enriched in UC rather than a generic pattern shared across all subjects.

**Fig. 2:**
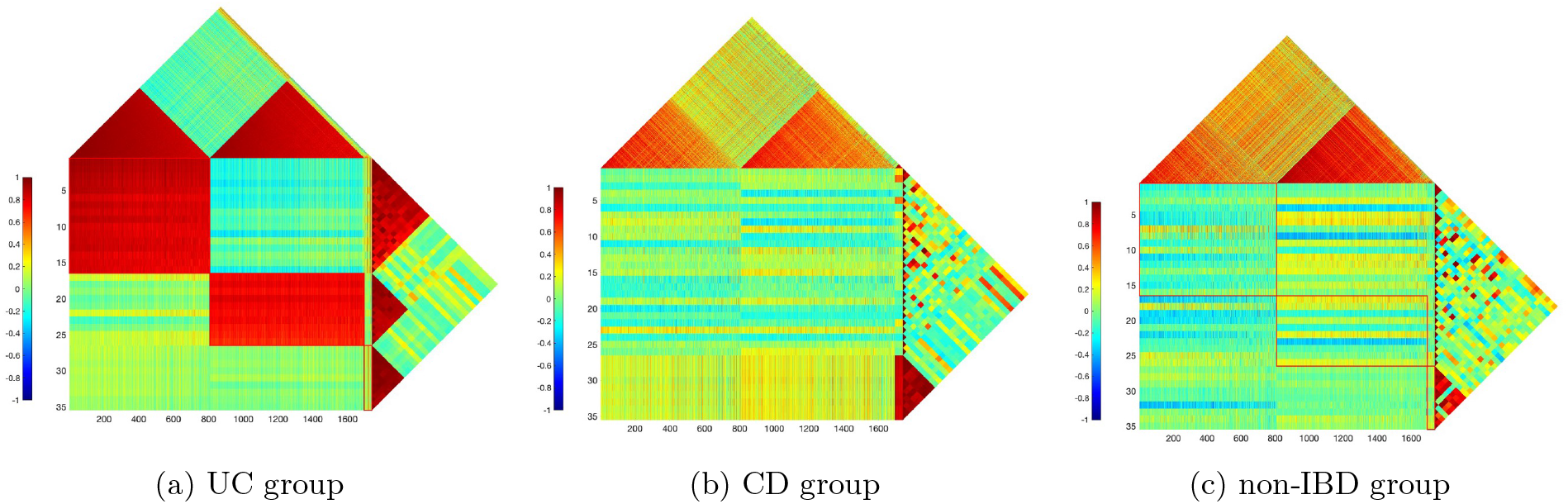
Recovered metabolite–transcript modules differ across IBD subgroups. For each group, the central panel shows the metabolite–transcript correlation matrix (Pearson correlation, color scale ranging from −1 to 1, with red indicating positive correlation and blue indicating negative correlation), whereas the upper and right triangular panels show the within-metabolite and within-transcript correlation structures, respectively. Variables are ordered identically across all three groups, and Modules 1, 2, and 3 appear sequentially along the main diagonal from upper left to lower right. The UC panel (a) shows the strongest and most coherent cross-omics blocks together with pronounced within-omics structure, whereas the corresponding patterns are weaker and more diffuse in CD (b) and non-IBD controls (c).

To quantify disease specificity more formally, we compared mean within-block Pearson correlation coefficients across disease groups and non-IBD controls (Table 2). Modules 1 and 2 showed substantially stronger mean cross-omics correlations in UC (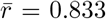 and 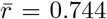, respectively) than in non-IBD controls (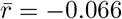 and 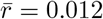), corresponding to effect sizes of 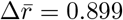 and 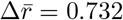. Module 3 showed the strongest enrichment in CD 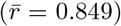 relative to controls 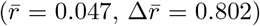, with near-zero cross-omics correlation in both UC and non-IBD groups. Permutation-based *P* values confirmed that all three between-group differences exceeded chance expectations (*p <* 10^−4^).

**Table 2:** Disease specificity of TriGer modules in the IBDMDB cohort. Module size is reported as the number of transcripts × number of metabolites selected by TriGer. Mean within-block Pearson correlation coefficients are shown for each detected module across CD, UC, and non-IBD controls. The effect size, 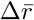 denotes the difference between the maximum mean within-block Pearson correlation observed across disease subgroups and the non-IBD control mean correlation, defined as 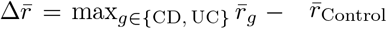. All between-group differences were statistically significant at *p <* 10^−4^ by permutation test.

| Module | Size | $\bar{r}_{CD}$ | $\bar{r}_{UC}$ | $\bar{r}_{Control}$ | Enriched subtype | $\Delta\bar{r}$ |
| --- | --- | --- | --- | --- | --- | --- |
| 1 | $929 \times 16$ | -0.001 | **0.833** | -0.066 | UC | 0.899 |
| 2 | $887 \times 10$ | -0.002 | **0.744** | 0.012 | UC | 0.732 |
| 3 | $45 \times 9$ | **0.849** | 0.142 | 0.047 | CD | 0.802 |

### Microbiome–metabolome coupling is attenuated in colorectal cancer

We then analyzed a colorectal cancer (CRC) dataset [22] integrating fecal microbiome and metabolome profiles. After restricting to samples with complete multi-omics measurements and focusing on the healthy-control versus CRC comparison, the final analysis included 127 healthy individuals and 123 patients with CRC. To retain associations that were strong in either condition, we combined group-specific evidence using the minimum *p*-value across the two groups (see Methods), thereby emphasizing condition-restricted signals rather than requiring concordance.

Fig. 3 displays the aggregated cross-omics association matrix between microbial genus and metabolite features. As in the IBD analysis, the original matrix shows no immediately apparent structure. After TriGer-guided reordering (Fig. 3, right panel; Step I), a prominent dense block emerges in the top-left corner, corresponding to a compact set of microbial genera and metabolites with strong associations in at least one condition. Fig. 4 shows the Step 2 refinement, in which row and column orderings are further guided by within-omics correlation blocks, yielding markedly more coherent structure across the cross-omics and within-omics matrices simultaneously.

**Fig. 3:**
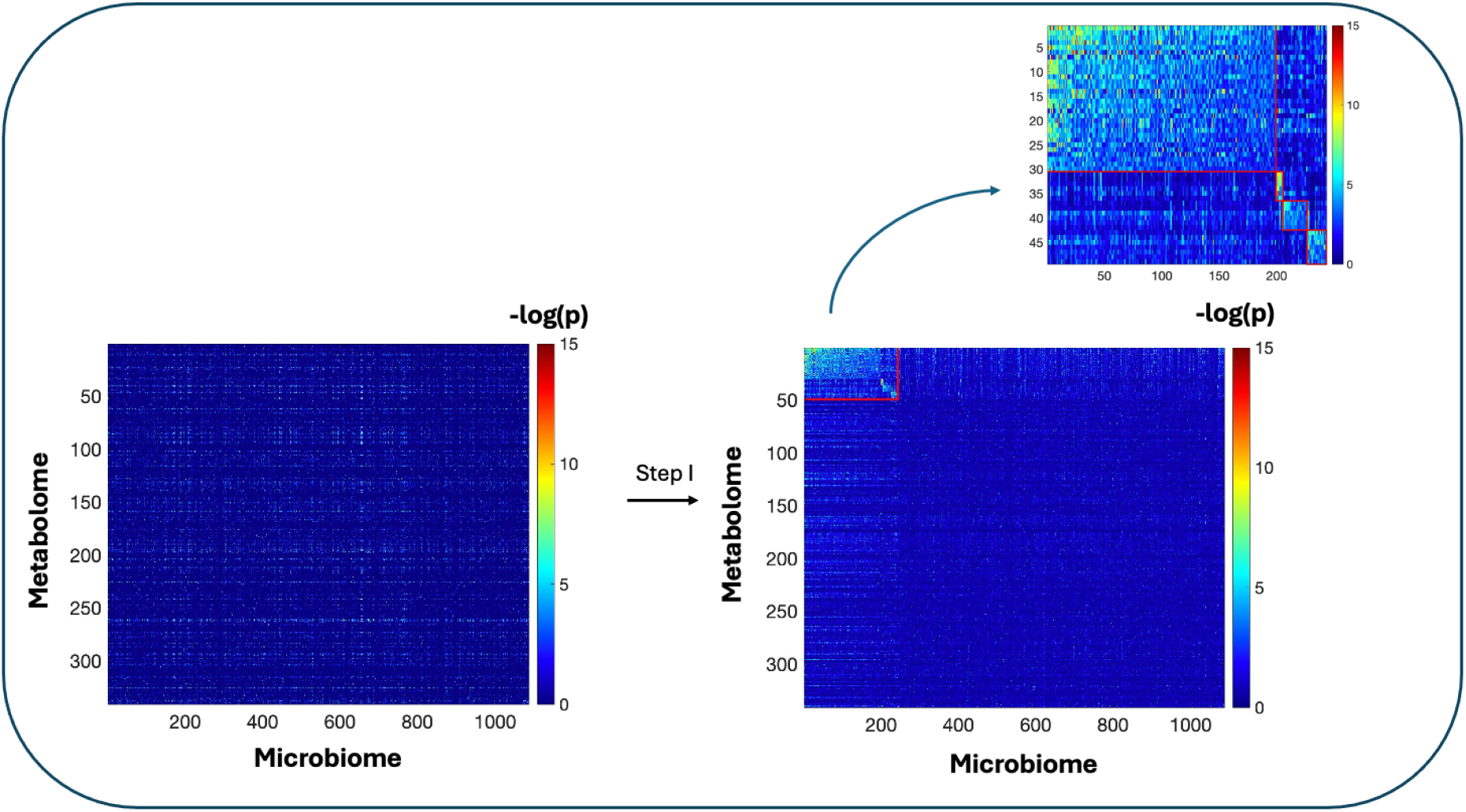
TriGer identifies a dense genus–metabolite association block in colorectal cancer data. Aggregated cross-omics association matrix between microbial genera (columns, *n* = 1,087) and metabolites (rows, *n* = 341), constructed from subgroup-specific correlations using a − log_10_(*p*) transformation. The left panel shows the original association matrix before reordering, and the right panel (Step I) shows the matrix after simultaneous reordering of rows and columns. The inset highlights the detected dense block (red box, top-left), corresponding to a compact set of microbial genera and metabolites with coordinated condition-specific associations. Warmer colors indicate stronger statistical significance.

**Fig. 4:**
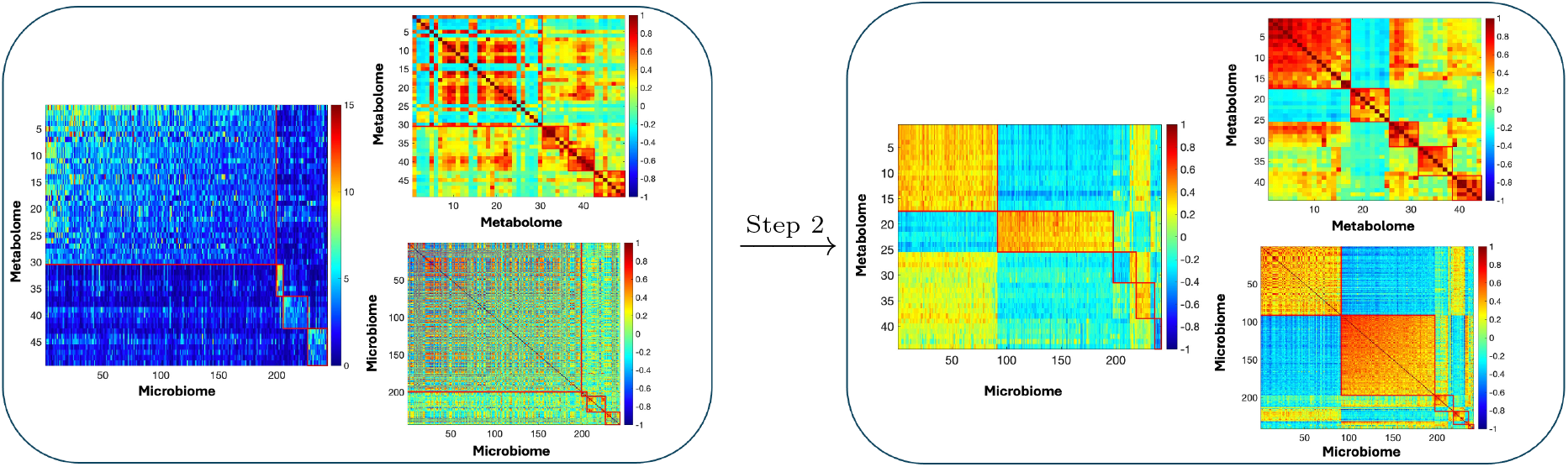
Within-omics refinement sharpens genus–metabolite module structure. The left panel shows the genus–metabolite association matrix (−log_10_(*p*)) together with the within-metabolite (upper right) and within-genus (lower right) Pearson correlation matrices before refinement. The right panel shows the corresponding matrices after Step 2 reordering, in which row and column order are guided primarily by within-omics correlation blocks. After refinement, compact genus and metabolite modules align more clearly with the cross-omics association blocks, yielding a more coherent structure across all three matrices. Color scales range from −1 to 1 for the within-omics correlation matrices and from 0 to 15 for the − log_10_(*p*) association matrix.

We next visualized the variables selected by TriGer in a joint matrix containing genus–genus correlations, metabolite–metabolite correlations, and genus–metabolite associations. Fig. 5 resolves five modules with aligned within- and cross-omics structure, indicating that the recovered features organize into coherent genus–metabolite modules rather than isolated associations. The contrast between healthy controls and CRC is most evident in Modules 3 and 5. In healthy controls, these regions show stronger and more sharply delimited cross-omics patterns, whereas in CRC the same regions are weaker and more diffuse. The results therefore point to selective erosion of structured genus–metabolite coupling in CRC rather than a uniform loss of association across the full matrix.

**Fig. 5:**
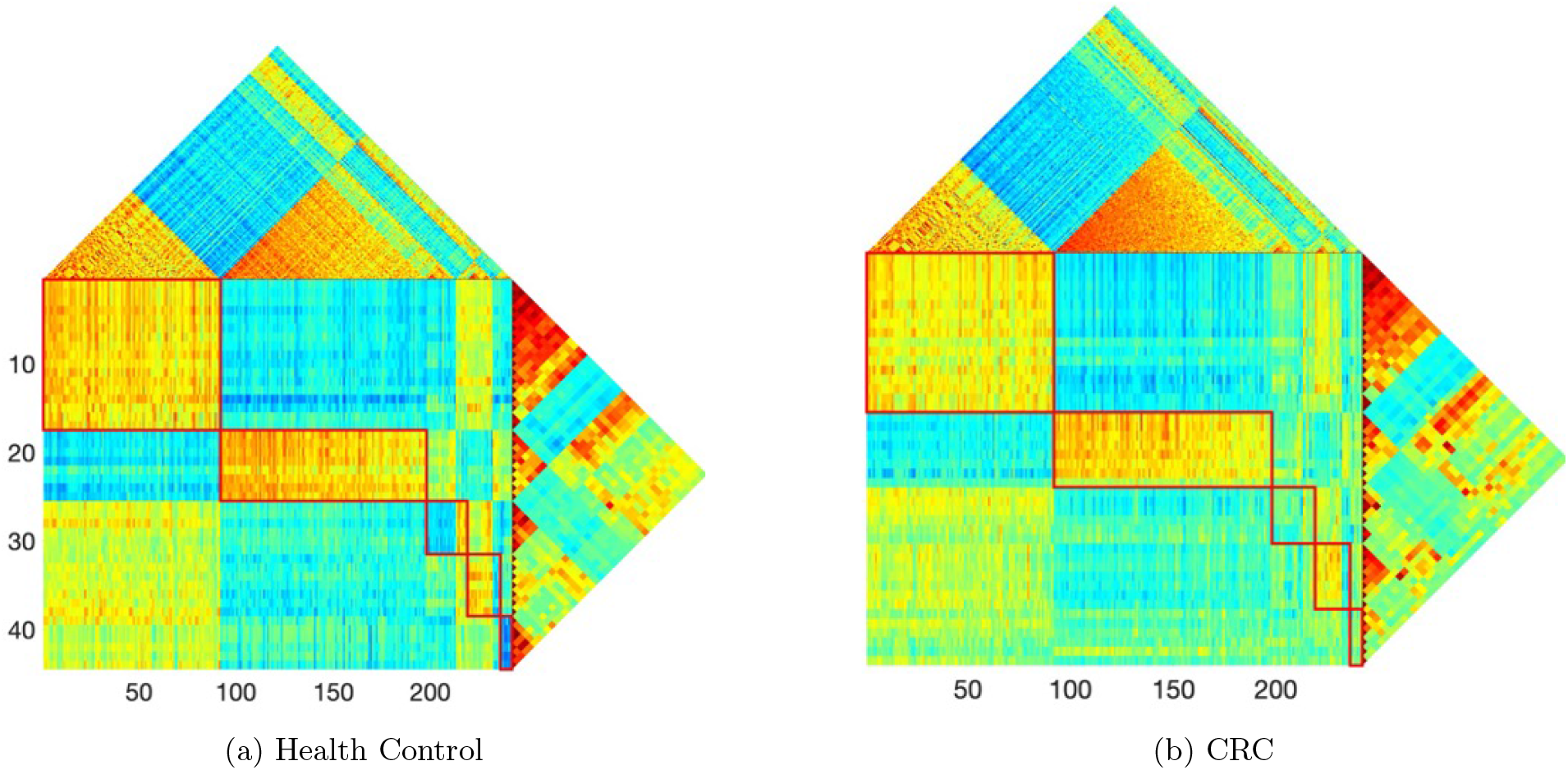
Genus–metabolite coupling is selectively attenuated in colorectal cancer. For each group, the central panel shows the genus–metabolite correlation matrix (Pearson correlation, color scale ranging from −1 to 1, with red indicating positive correlation and blue indicating negative correlation), whereas the upper and right triangular panels show the within-metabolite and within-genus correlation structures, respectively. Red boxes delineate the five detected modules, which are ordered sequentially along the main diagonal from upper left (Module 1) to lower right (Module 5). Modules 3 and 5 showed the most pronounced between-group differences and are the focus of the main text. They are more strongly organized in healthy controls (a) than in CRC (b), where the corresponding blocks are weaker and more diffuse, consistent with selective disruption of coordinated genus–metabolite structure in cancer.

To assess the robustness of the detected modules to sampling variability, we performed a resampling analysis in which 100 individuals were randomly drawn without replacement from each group (healthy controls: *n* = 127; CRC: *n* = 123) across *B* = 100 iterations. In each iteration, group-specific correlation matrices were recomputed from the subsampled data, the aggregated association matrix was reconstructed, and TriGer was reapplied under identical parameter settings. Module stability was quantified by the recall of the originally identified features, defined as the proportion of features from the full-data solution recovered in each subsampled run. Across 100 iterations, mean recall was 82.0% for metabolite features and 78.0% for microbiome features. Given that each subsample retained approximately 80% of the original observations, these recall rates indicate that the core module structure is consistently recoverable from reduced data, supporting the stability of the detected modules under sampling perturbation.

## Discussion

### Summary

Multi-omics studies are increasingly expected to move beyond lists of pairwise associations and toward representations that better reflect how biological systems are organized. That transition remains difficult in practice because cross-omics signals are often weak, high dimensional, and distributed across groups of features rather than concentrated in a few dominant pairs. Existing methods address parts of this problem, but they typically emphasize either global latent structure or pairwise cross-layer association, without explicitly preserving within-omics coherence and cross-omics coupling at the same time. As a result, biologically meaningful signals can appear either too diffuse to interpret or too unstable to reproduce.

TriGer was designed to address that gap by treating the target of inference as a structured module rather than as an isolated association. By jointly prioritizing dense cross-omics blocks and coherent within-omics organization, the method recovers modules that are both statistically compact and biologically interpretable. This distinction mattered in the simulation studies: TriGer was the only approach that consistently balanced sensitivity and specificity across the primary planted blocks and the nested blocks, whereas the competing methods showed characteristic failure modes. MOFA and JIVE tended to miss weaker embedded signal, FABIA often recovered signal at the cost of over-inclusion, and sCCA-PMA and DIABLO were more vulnerable to contamination by false positives or instability across settings. Taken together, these patterns suggest that explicitly modeling block structure is advantageous when the true signal is modular rather than diffuse.

The two real-data applications illustrate the practical value of that shift in emphasis. In the IBD cohort, the method resolved disease-stratified metabolite–transcript modules, including UC-associated lipid and mitochondrial signatures and a CD-associated lysophospholipid–myeloid inflammatory axis. In the CRC application, it identified microbiome–metabolome modules that were structured in healthy controls but selectively attenuated in cancer, pointing to a loss of coordinated ecological-metabolic organization rather than a uniform weakening of all associations. Viewed together, these findings support the idea that module-level structure can serve as a useful intermediate scale between single features and global latent factors, enabling mechanistic interpretation while remaining computationally tractable in high-dimensional data.

### Cross-modality clustering reveals disease-specific host–metabolite modules in inflammatory bowel disease

The IBD application illustrates why a module-based representation can be more informative than an exhaustive correlation map. Rather than returning thousands of metabolite–transcript pairs that are difficult to compare across disease groups, TriGer resolved a small number of coordinated cross-modality units that could be interpreted at the level of pathways and disease mechanisms. In turn, this made it easier to separate patterns shared across groups from those enriched in UC or CD, and to connect the observed metabolite structure to host transcriptional programs in a biologically coherent way.

As shown in Fig.2, in UC, TriGer identified a lipid-dominant fecal module (Module 1) absent in CD and non-IBD controls, composed of long-chain acylcarnitines (C14-, C18:1-, C18:2-carnitine), diverse fatty acids (myristate/myristoleate, phytanate, heptadecanoate, eicosenoate/eicosadienoate), N-acylethanolamides (palmitoylethanolamide, linoleoyl ethanolamide), very-long-chain ceramides (Cer(d18:1/22:0/24:0/24:1)), and oxalate. This constellation aligns with UC fecal metabolomics reporting inflammation-associated elevations in ceramides, long-chain fatty acids, and acylcarnitines,[23] and with mucosal lipidomics identifying Cer(d18:1/24:0) as discriminatory between active UC, remission, and healthy controls.[24] Importantly, the module was not only a within-metabolome pattern: it exhibited UC-specific cross-modality coupling to host gene programs controlling mitochondrial bioenergetics (oxidative phosphorylation, respiratory electron transport, Complex I–IV assembly, ATP synthesis) and oxidative-stress defense (hydrogen peroxide/ROS detoxification). This joint structure supports a model in which UC mucosal/epithelial mitochondrial stress and remodeling shapes luminal lipid metabolism, consistent with emerging syntheses linking epithelial mitochondrial dysfunction and altered oxidative phosphorylation to intestinal inflammation and barrier disruption.[25, 26] In mechanistic terms, long-chain acylcarnitines accumulate when mitochondrial fatty-acid oxidation is incomplete,[27] consistent with the “energy deficiency” model of UC colonocytes that emphasizes impaired oxidative metabolism.[28, 29] Concomitantly, very-long-chain ceramides can directly perturb respiratory chain activity while promoting reactive oxygen species (ROS) generation,[30] offering a plausible link between sphingolipid remodeling and the observed oxidative-stress gene signatures. Together, these findings highlight a UC-specific axis in which mitochondrial energetic insufficiency and ceramide-associated lipotoxic stress co-vary with fecal lipid remodeling, nominating a coherent, testable integrative mechanism rather than isolated metabolite changes.

A second UC-restricted module (Module 2) implicated a distinct but complementary biology at the interface of microbial metabolism, nutrient stress signaling, and cellular quality control. This module included microbiome-linked amino-acid catabolites spanning tryptophan and aromatic amino-acid metabolism (indole-3-acetic acid and p-hydroxyphenylacetate) and histidine/imidazole products (imidazolelactate, imidazoleacetic acid), pyrimidine degradation intermediates (N-carbamoyl-*β*-alanine and *β*-alanine), and lipid stress mediators (*α*-linolenate, 9,10-DiHOME, and short-chain ceramide). The UC-specific correlation of these metabolites was mirrored by selective coupling to host pathways governing proteostasis and RNA fate (ubiquitin–proteasome, autophagy/lysosome organization, P-body assembly, regulation of mRNA stability), as well as upstream nutrient/stress control nodes (mTOR/FoxO). Several components of the module have independent links to colitis biology: indole-3-acetic acid is a gut bacteria–associated tryptophan metabolite[31] reported to be decreased in UC feces with experimental evidence of colitis protection upon supplementation,[32] while p-hydroxyphenylacetate is a major colonic microbiota–derived aromatic metabolite with host bioactivity and has been implicated as anti-inflammatory/barrier-supportive in colitis models.[33, 34] On the lipid side, 9,10-DiHOME is generated downstream of soluble epoxide hydrolase activity,[35] and soluble epoxide hydrolase deficiency/inhibition suppresses experimental colitis and related inflammatory pathology.[36, 37] Finally, ceramide provides a mechanistic bridge from lipid stress to the observed phosphorylation/dephosphorylation and protein-turnover programs, as ceramide can activate PP2A,[38] impair mitochondrial respiratory chain function,[39] and modulate mTOR-controlled autophagy.[40] Collectively, this module supports a UC-specific microbial metabolite–lipid stress–proteostasis axis, suggesting that epithelial/immune stress adaptation (UPS–autophagy cross-talk and post-transcriptional regulation) is not merely a downstream consequence of inflammation but a coordinated component of UC-specific host–microbial metabolic coupling.

In contrast, the CD-specific cross-modality module (Module 3) was anchored by lysophospholipid and ether-phospholipid remodeling, comprising multiple lysophosphatidylcholines (LPCs; including LysoPC 20:4) and lysophosphatidylethanolamides (LPEs), together with plasmalogen PC and plasmalogen LPC species. LPC is produced from phosphatidylcholine by phospholipase activity and functions as a bioactive lysolipid that signals through GPCRs (notably G2A/GPR132), promoting immune cell migration and inflammatory signaling.[41–43] This biochemical signature aligned with the coupled gene programs enriched for leukocyte chemotaxis/chemokine signaling, cytokine production, Fc*γ* receptor signaling, respiratory burst/oxidoreductase activity, calcium homeostasis, and Toll-like receptor signaling, consistent with activated myeloid programs in inflamed tissue, because LPC can act as a chemotactic cue via G2A and engage TLR2/TLR4-linked signaling in myeloid cells.[42–44] The presence of plasmalogen PCs/LPCs further supports an immune-redox and lipid-mediator context: plasmalogens have been proposed to participate in redox balance,[45] and their hydrolysis can mobilize polyunsaturated fatty acids that feed downstream inflammatory lipid mediator pathways.[46] Consistent with this interpretation, CD-focused fecal metabolomics has reported elevation of specific LysoPC species in CD compared with healthy controls, reinforcing the disease relevance of lysophospholipid accumulation in CD-associated immune activation.[47] Notably, these CD modules differed qualitatively from UC modules: rather than primarily reflecting mitochondrial energetic failure, CD modules emphasized membrane lipid remodeling coupled to chemotaxis/complement/Fc signaling and oxidative burst programs, supporting mechanistic heterogeneity between the two diseases that becomes clearer when metabolite and gene modules are learned jointly.

More broadly, these IBD results show how cross-modality clustering can sharpen biological interpretation in heterogeneous human cohorts. First, the modules compress high-dimensional association structure into a small number of interpretable units. Second, they support mechanistic framing at the level of processes rather than individual markers, which is more useful for cross-study comparison and follow-up experimentation. Third, disease-restricted coupling patterns, such as modules enriched in UC but not CD or controls, may help prioritize context-dependent biomarkers and pathways that would be diluted in pooled analyses.

### Colorectal cancer disrupts structured microbe–metabolite network organization

The CRC application highlights a complementary use case: identifying not only which variables cluster together, but also where structured cross-omics organization breaks down in disease. In this setting, the most striking signal was not the appearance of entirely new cancer-specific modules, but the attenuation of genus–metabolite modules that were well organized in healthy controls. That pattern is consistent with disease-associated disruption of homeostatic metabolic cross-feeding and broader ecological control within the gut environment.

In the first module (Fig.5, Module 3), healthy individuals exhibited a cross-modality antagonism between amino-acid–linked hydroxy acids and Lachnospiraceae genera. The metabolite set is chemically and biogenically coherent, spanning aromatic and branched-chain hydroxy acids, 3-phenyllactate (a microbial reduction product of phenylalanine metabolism, commonly produced by lactic acid bacteria),[48] 2-hydroxy-4-methylpentanoate (2-hydroxyisocaproate/leucic acid; a leucine catabolite produced in humans and some microorganisms),[49] and (hydroxy)phenyl-propionate–like metabolites that are typical gut microbial products of aromatic substrates[50], together with proline/hydroxyproline-linked features. These metabolites were negatively associated with multiple Lachnospiraceae genera (e.g., *Agathobacter, Anaerobutyricum, Dorea, Lachnospira*, and *Falcatimonas*). Lachnospiraceae are prominent anaerobic fermenters that can metabolize host-derived amino acids, aromatic compounds, and mucin glycoprotein breakdown products,[51–54] and many members (including *Agathobacter, Anaerobutyricum, Dorea*) are linked to SCFA production and butyrate-associated community states.[55–57] A parsimonious ecological interpretation is that, in the healthy gut, these taxa mark (or contribute to) a fermentation regime that limits net accumulation of amino-acid–derived hydroxy acids—through direct utilization/conversion and/or via maintenance of carbohydrate-fermentative, SCFA-producing niches that suppress proteolytic/aromatic fermentation end-products.[58, 59] Strikingly, this antagonistic coupling was lost in CRC, indicating disease-associated decoupling between microbial community structure and amino-acid–linked hydroxy acids, consistent with broader CRC frameworks implicating altered microbial metabolism, including shifts in amino-acid metabolism and fermentation patterns, in tumorigenesis.[60, 61] Importantly, the module also provides a mechanistic anchor for host–tumor interaction. Proline/hydroxyproline–linked features may reflect altered extracellular matrix turnover and collagen catabolism, and growing evidence implicates proline metabolism in malignant cell programs and in shaping the tumor microenvironment. [62, 63]. Hydroxyproline, in particular, has been proposed to support pro-survival phenotypes in cancer cells[64] and has been reported to potentiate IFN-*γ*–induced PD-L1 expression while inhibiting autophagic flux,[65, 66] linking this axis to immune-evasion and stress-adaptation programs. The aromatic branch includes 3-phenyllactate (PLA), which has been associated with tumor-relevant bioactivity: PLA can promote migration and invasion via MMP-9 signaling in cervical cancer cells[67] and suppress apoptosis in prostate cancer cells, potentially through increased expression of cysteine desulfurase (NFS1), alongside reductions in TNF-*α*.[68] In this context, the loss of the healthy antagonistic coupling may permit increased accumulation of metabolites with potential tumor-relevant activity, providing a plausible route by which ecological disruption translates into altered luminal exposures that impinge on epithelial signaling, redox balance, and inflammation.

Module 5 captured a distinct facet of CRC-associated rewiring: a healthy-specific, internally coherent metabolite consortium with coordinated negative coupling to SCFA-linked commensals. In healthy controls, this module united metabolites that map to cancer-relevant metabolic axes: redox buffering and glutathione turnover (5-oxoproline/pyroglutamate, a *γ*-glutamyl cycle intermediate [69] that has been used as a marker of glutathione depletion/oxidative stress [70, 71]), branched-chain and lysine amino-acid catabolism (4-methyl-2-oxopentanoate/*α*-ketoisocaproate from leucine catabolism [72, 73] and saccharopine from the mitochondrial lysine degradation pathway [74]), and host–microbe immunometabolic signaling (agmatine (a microbiota-derivable arginine decarboxylation product) [75], itaconate (an ACOD1/IRG1-linked immunometabolite with antimicrobial/immunoregulatory roles) [76–78]). Agmatine is particularly notable in CRC context because a recent mechanistic study implicated commensal microbiota–derived agmatine in inflammation-associated colorectal tumorigenesis via activation of Wnt signaling, [79] while itaconate, classically an ACOD1/IRG1-linked macrophage immunometabolite, has emerging, context-dependent roles in cancer immunity, including reports that itaconate/IRG1 signaling can shape tumor immune escape and response to immune checkpoint therapy. [80–82] Finally, the inclusion of the amino sugar D-mannosamine connects the module to hexosamine/sialic-acid–linked glycocalyx biology, relevant because tumor-associated sialylation is widely implicated in cancer progression and immune evasion through Siglec–sialic acid interactions. [83–85] Among healthy controls only, this metabolite consortium was negatively correlated with commensal anaerobes from Ruminococcaceae/Oscillospiraceae/Butyricicoccaceae (e.g., Agathobaculum, Lawsonibacter, Faecalibacterium), which are core ecosystem members frequently associated with SCFA production (notably butyrate).[86–88] Given the roles of SCFAs in epithelial energy supply, anti-inflammatory signaling, and barrier integrity,[89–91] one interpretation is that SCFA-linked community states are associated with lower net accumulation of nitrogen/redox and amino-acid catabolic intermediates, either via direct utilization/cross-feeding or indirectly via improved mucosal redox homeostasis, consistent with the central role of oxidative stress and metabolic reprogramming in CRC. [92, 93]

Taken together, the CRC results reinforce the value of using cross-modality modules as the primary unit of interpretation. The recovered modules connect taxonomic community states, including Lachnospiraceae- and SCFA-associated commensals, to chemically coherent metabolite programs involving amino-acid catabolism, redox buffering, immunometabolic signaling, and hexosamine-related features. Just as importantly, they indicate that CRC is characterized by selective loss of structured coupling rather than by a uniform collapse of all genus–metabolite associations. In practical terms, these modules offer testable candidates for disease stratification, targeted validation of specific metabolite–genus axes, and future intervention strategies aimed at restoring cross-feeding and immune-metabolic balance.

## Conclusions

TriGer addresses the challenge of recovering structured cross-omics modules that are simultaneously dense across molecular layers and internally coherent within each layer. By shifting the unit of inference from isolated feature pairs to coordinated module-to-module interactions, it aggregates weak but concordant signals into compact, interpretable units while preserving the internal organization of each layer. Our benchmarking demonstrates that this design substantially improves upon dimension-reduction and association-based methods in both sensitivity and specificity, and on real data it resolved disease-stratified metabolite–transcript modules in inflammatory bowel disease and revealed selective attenuation of genus– metabolite coupling in colorectal cancer. Because the framework extends naturally to count-valued omics through a Gaussian copula formulation and accommodates one-to-multiple module mappings, it offers a practical route to characterizing system-level molecular organization and linking coordinated multi-omic programs to complex disease phenotypes.

## Methods

### Method overview and theoretical details

TriGer is designed to identify structured cross-omics modules that are simultaneously dense across two molecular layers and internally coherent within each layer. The method takes as input a cross-omics association matrix together with two within-omics correlation matrices. It then searches for pairs of feature subsets that define a dense bipartite block in the cross-omics matrix while remaining densely connected in the two within-omics graphs. In this way, the recovered module is required to satisfy both cross-layer relevance and within-layer coherence.

At a high level, the procedure has two stages. Step I identifies a dense cross-omics block by iteratively pruning weakly connected nodes from the bipartite association graph. Step II refines the retained features using the within-omics graphs so that the final module also exhibits strong internal correlation structure in each modality. This design places primary emphasis on cross-omics association while using within-omics structure as a supporting constraint. The formal objective function, optimization details, and supporting theoretical arguments are given below.

Fig. 6 illustrates these targets from a graph-theoretic perspective. The upper panel shows the desired configuration: a bipartite subgraph with dense cross-omics associations together with strong within-omics connectivity among the selected features in each layer. The lower panel shows the complementary case, in which within-omics coherence is present but cross-omics association is absent. This latter pattern lies outside the recovery target of TriGer and highlights a central design principle of the method: within-layer co-variation alone is not sufficient to define a biologically meaningful cross-omics module.

**Fig. 6:**
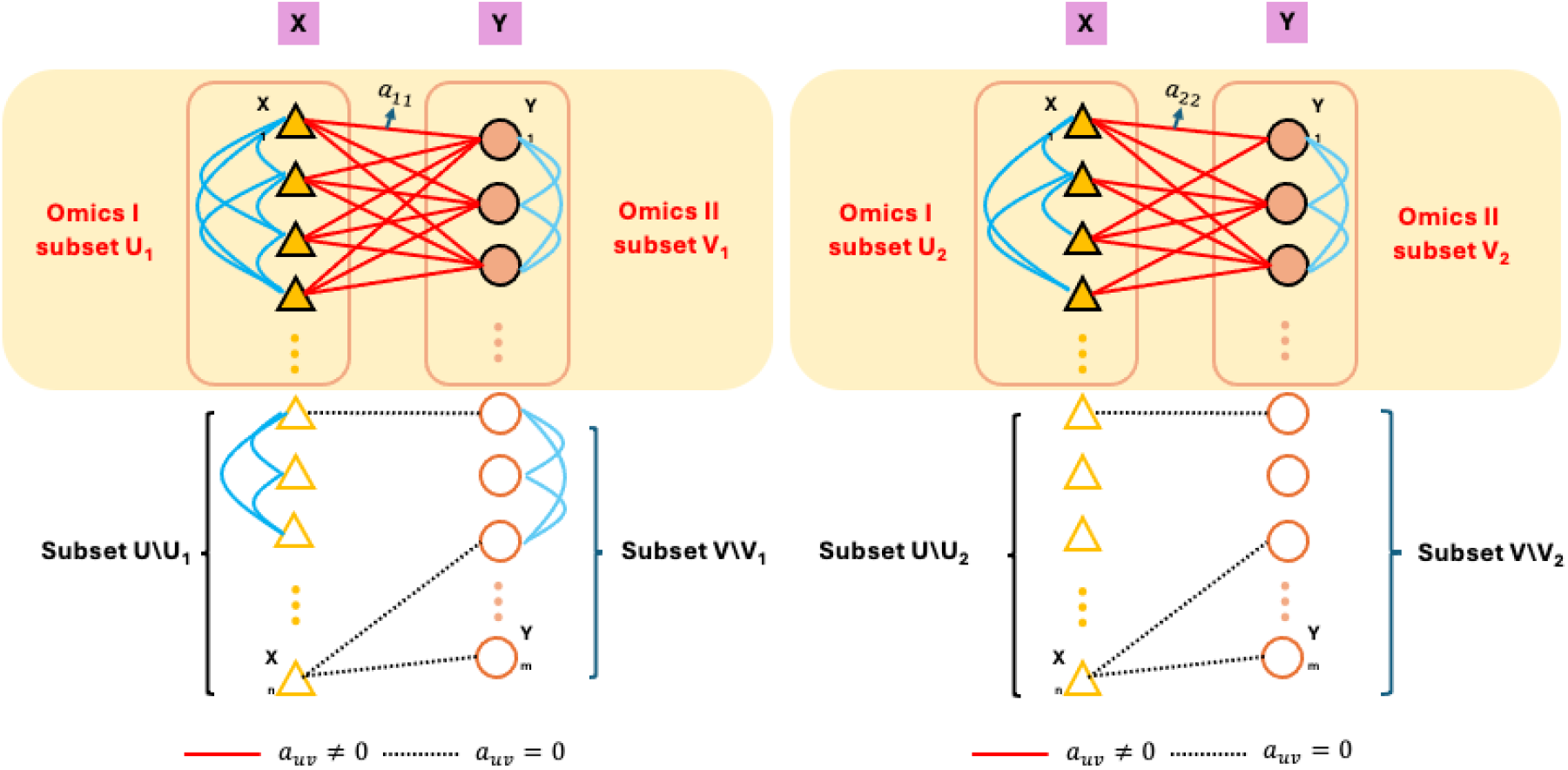
Graph-theoretic targets of TriGer. The yellow region in the left diagram represents the primary target: a dense cross-omics subgraph whose corresponding feature sets are also strongly connected within their respective omics layers. The yellow region in the right diagram illustrates a configuration in which cross-omics connectivity remains strong but within-omics coherence is weaker, representing a less preferred but still recoverable pattern. The unshaded region denotes strong within-omics structure that does not extend across layers and is therefore not the main target of extraction. Solid lines denote nonzero cross-omics associations (*a*_*uv*_ ≠ 0) and dotted lines denote absent associations (*a*_*uv*_ = 0).

Fig. 7 shows the same idea in matrix form using simulated data. The left panels show the Omics I (500 × 60) and Omics II (500 × 80) data matrices. The center-left column displays the three input matrices (Σ_*XX*_, Σ_*XY*_, and Σ_*Y Y*_) before reordering, in which no apparent block structure is visible. The center-right column shows the same three matrices after TriGer module detection and reordering; coherent block structure emerges simultaneously across all three matrices, confirming that each recovered module is supported jointly by cross-layer coupling and within-layer coherence. The right panel combines the three reordered matrices into a unified diamond display, with Σ_*XY*_ at the center flanked by Σ_*XX*_ (right triangle) and Σ_*Y Y*_ (upper triangle). The red boxes mark detected modules, each exhibiting strong cross-omics association together with coherent within-omics structure. Features with strong within-omics correlations but no corresponding cross-omics signal are not selected, as illustrated in Fig. 7.

**Fig. 7:**
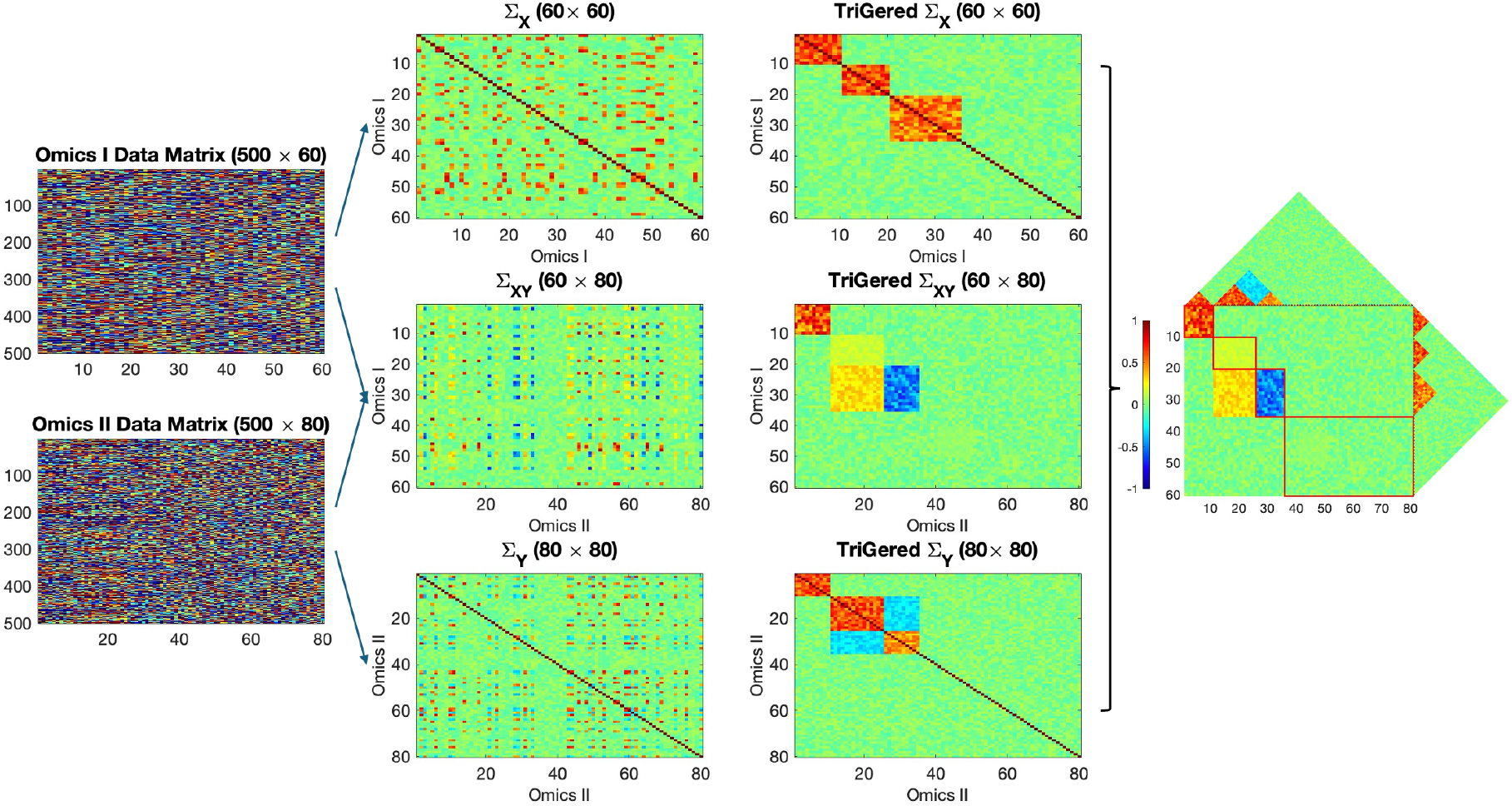
TriGer triple-graph representation and module detection. The left panels show the Omics I (500 × 60) and Omics II (500 × 80) data matrices. The center-left column displays the three input matrices (within-Omics I correlations Σ_*X*_, 60 × 60; cross-omics associations Σ_*XY*_, 60 × 80; within-Omics II correlations Σ_*Y*_, 80 × 80) before reordering, with no apparent block structure. The center-right column shows the same three matrices after TriGer module detection and reordering, where coherent block structure emerges simultaneously across all three matrices. The right panel combines the reordered matrices into a unified diamond display, with Σ_*XY*_ at the center flanked by Σ_*X*_ (right triangle) and Σ_*Y*_ (upper triangle). Red boxes mark the detected modules, illustrating that TriGer recovers structured cross-omics modules supported jointly by cross-layer coupling and within-layer coherence.

### Simulation design and benchmarking protocol

We evaluated TriGer under two simulation settings designed to mimic block-structured cross-omics signal with nested blocks. Scenario 1 contained 450 Omics I features and 550 Omics II features and included one primary planted block together with one smaller nested block. The primary block comprised 40 Omics I features and 60 Omics II features with stronger signal, whereas the nested block comprised 10 Omics I features contained within the primary Omics I set and an additional 20 Omics II features with a weaker but still structured signal.

Scenario 2 was more challenging and included 400 Omics I features and 600 Omics II features. In this setting, the simulated cross-omics matrix contained one primary block and two overlapping nested blocks. The primary block consisted of 30 Omics I features and 50 Omics II features. Nested block 1 consisted of 20 Omics I features drawn from the primary Omics I set together with 30 Omics II features. Nested block 2 consisted of 15 Omics I features and 10 Omics II features embedded within nested block 1. Across scenarios, signal strength and block configuration were varied to evaluate recovery of both the main planted structure and the more difficult embedded substructures.

Benchmarking compared TriGer with representative dimension-reduction and association-based methods, including MOFA, JIVE, FABIA, sCCA-PMA, and DIABLO. Performance was summarized using true positive rate (TPR) and true negative rate (TNR) for each planted block, thereby assessing both sensitivity to true signal and resistance to false positive inclusion.

### Cohort preprocessing and construction of association matrices

For each application dataset, preprocessing was first carried out separately within each omics layer, followed by construction of subgroup-specific association matrices and within-omics correlation matrices. These matrices served as the direct input to TriGer.

#### IBD cohort

The IBD analysis used matched metabolomics and transcriptomics profiles from the Inflammatory Bowel Disease Multiomics Database (IBDMDB) [8]. After restricting to participants with both data types available, the final paired dataset contained 76 individuals: 32 with Crohn’s disease (CD), 22 with ulcerative colitis (UC), and 22 non-IBD controls. Within each subgroup, pairwise metabolite–transcript association tests were performed to obtain subgroup-specific *p*-value matrices. To retain subgroup-restricted signals while avoiding the requirement that the same association be present in all three groups, we defined an aggregated screening matrix by taking the strongest transformed association signal across the three subgroups:

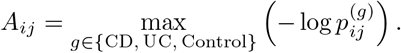

The selected modules were then projected back to the subgroup-specific matrices to evaluate whether the corresponding structure was shared or disease-specific.

#### CRC cohort

The CRC analysis used the public data set [22] that integrates the fecal microbiome and metabolome profiles throughout the progression of colorectal cancer. Samples with complete measurements across the two omics layers were retained. Metabolites with fewer than 5% nonzero observations were removed, and low-prevalence microbial genera with nonzero abundance in fewer than 10% of samples were excluded. A compositional bias-correction procedure was then applied to the microbiome data [94]. For the case– control analysis, intermediate groups were excluded and the final comparison focused on 127 healthy individuals and 123 CRC cases. Group-specific cross-omics association tests were performed separately in controls and CRC, and the resulting evidence was combined by taking the maximum of −*log*(*p*) value across the two groups so that strong condition-specific signals were retained in the screening matrix.

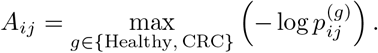

### Data structure and problem set up

We observe continuous Omics I and Omics II measurements from *S* independent samples. Let **x**^(*i*)^ ∈ ℝ^*m*^ and **y**^(*i*)^ ∈ ℝ^*n*^ denote the feature vectors, with index sets *U* (|*U*| = *m*) and *V* (|*V*| = *n*). Define the joint observation vector **s**^(*i*)^ = (**x**^(*i*)⊤^, **y**^(*i*)⊤^)^⊤^ ∈ ℝ^*m*+*n*^, and let **z**^(*i*)^ ∈ ℝ^*q*^ denote a vector of sample-level covariates (e.g., age, sex, and batch effects) included to adjust for confounding in the marginal association tests. The conditional joint covariance matrix of **s**^(*i*)^ given **z**^(*i*)^ is then

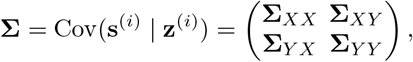

where **Σ**_*XX*_ ∈ ℝ^*m*×*m*^ and **Σ**_*Y Y*_ ∈ ℝ^*n*×*n*^ encode within-layer covariance structure, and **Σ**_*XY*_ ∈ ℝ^*m*×*n*^ encodes cross-layer associations [95]. The cross-layer block **Σ**_*XY*_ naturally defines a weighted bipartite graph between Omics I and Omics II features and is the primary target of TriGer inference. We note that the conditional approach, encoding cross-layer association through the off-diagonal block of the precision matrix **Ω** = **Σ**^−1^, is not appropriate here. By the Schur complement formula for block matrix inversion,

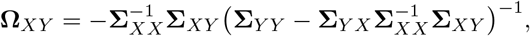

which mixes all three blocks **Σ**_*XX*_, **Σ**_*XY*_, and **Σ**_*Y Y*_. As a result, block structure in **Σ**_*XY*_ is not preserved in **Ω**_*XY*_ after inversion, and clear block patterns observed in the marginal covariance matrix are lost in the conditional representation [96], motivating a covariance-domain approach for recovering the dense bipartite block patterns that TriGer targets. Furthermore, in the high-dimensional setting where *m* +*n* ≫ *S*, the precision matrix **Ω** is not consistently estimable without strong sparsity assumptions, whereas marginal pairwise associations remain well-posed for each (*u, v*) pair independently [97]. We therefore estimate associations marginally for each pair (*u, v*) with *u* ∈ *U* and *v* ∈ *V*, where *β*_*uv*_ is the marginal association coefficient between Omics I feature *u* and Omics II feature *v*, conditional on covariates **z**^(*i*)^. The matrix ***β*** = {*β*_*uv*_} ∈ ℝ^*m*×*n*^ collects all conditional marginal association coefficients and serves as an estimator of the cross-layer block **Σ**_*XY*_, preserving its bipartite structure. The corresponding edge set *A* = {*a*_*uv*_} with *a*_*uv*_ = **1**(*β*_*uv*_ ≠ 0) defines a weighted bipartite graph *B* = (*U, V, A*) between Omics I and Omics II features.

Although the framework is presented for continuous measurements, it extends directly to count-valued omics data (e.g. microbiome feature tables, single-cell RNA-seq) via a Gaussian copula substitution, and together continuous and count representations cover virtually most of standard omics data types.

### Multivariate-to-multivariate inference from a graph perspective

Naive enumeration of all *mn* pairwise hypotheses 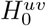 treats each Omics I–Omics II feature pair in isolation, imposing a multiple-testing burden that grows with the product of the two feature dimensions and discarding the coordinated structure inherent to biological systems. For example, a co-regulated metabolite set may jointly associate with a co-expressed gene module, a pattern that is effectively invisible to pairwise testing. Specifically, for each pair (*u, v*) with *u* ∈ *U* and *v* ∈ *V*, the marginal hypothesis is

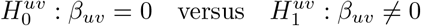

We therefore represent the association matrix ***β*** ∈ ℝ^*m*×*n*^ as a bipartite graph *B* = (*U, V, A*) as defined above; recovering the support of ***β*** is thus equivalent to recovering the edge set *A*. Inference is consequently redirected toward identifying dense induced subgraphs *B*[*U*_*c*_, *V*_*d*_], each encoding a coherent cluster-to-cluster association between a subset of Omics I features *U*_*c*_ ⊂ *U* and a subset of Omics II features *V*_*d*_ ⊂ *V*. This reformulation compresses a high-dimensional testing problem into a smaller number of biologically interpretable module-level hypotheses. In high-dimensional applications the graph is typically sparse, with only a small fraction of all possible cross-omics pairs carrying meaningful signal.A candidate cross-omics module is therefore represented by an induced bipartite subgraph *B*[*U*_*c*_, *V*_*d*_] = (*U*_*c*_, *V*_*d*_, *A*[*U*_*c*_, *V*_*d*_]), where *U*_*c*_ ⊂ *U* and *V*_*d*_ ⊂ *V*, and *A*[*U*_*c*_, *V*_*d*_] = {(*u, v*) : *u* ∈ *U*_*c*_, *v* ∈ *V*_*d*_, *β*_*uv*_ 0} denotes the set of active edges within the candidate module. The basic structural assumption is that the average edge density inside the target subgraph exceeds the corresponding background density outside the subgraph:

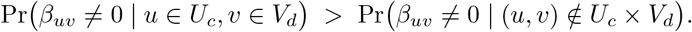

Or equivalently,

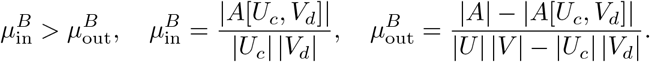

To incorporate structure within each modality, we also estimate two weighted within-omics graphs: *G* = (*U, E, W* ^(1)^) for Omics I and *H* = (*V, F, W* ^(2)^) for Omics II, where *E* and *F* encode within-layer associations such as correlations or partial correlations, and *E*[*U*_*c*_] = {(*u, u*^*′*^) ∈ *E* : *u, u*^*′*^ ∈ *U*_*c*_} and *F* [*V*_*d*_] = {(*v, v*^*′*^) ∈ *F* : *v, v*^*′*^ ∈ *V*_*d*_} denote the active edges within each induced subgraph. For notational convenience, |*U*_*c*_|^2^ and |*V*_*d*_|^2^ are used as approximations to the number of possible edges within each induced within-omics subgraph, treating the within-omics graphs as directed for uniformity. For a candidate module (*U*_*c*_, *V*_*d*_), the induced subgraphs *G*[*U*_*c*_] and *H*[*V*_*d*_] are expected to be denser than background if the selected features form coherent within-omics modules as well as a dense cross-omics block. For Omics I, this can be written as

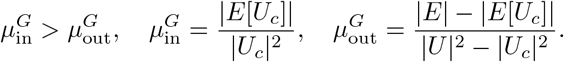

Analogously, for Omics II we require

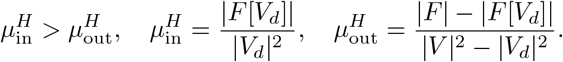

In this notation, the superscripts *B, G*, and *H* distinguish the bipartite cross-omics graph and the two within-omics graphs, respectively, while the subscripts “in” and “out” denote average density inside and outside the candidate module. The target pattern is therefore not merely a collection of cross-omics edges, but a coordinated structure that is dense across all three graphs.

Even when individual associations *β*_*uv*_ are statistically significant, isolated pairs are not the primary target of our inference. A single significant *β*_*uv*_ may reflect noise or a feature-specific effect rather than a meaningful biological relationship. We focus instead on module-to-module associations: sets *U*_*c*_ ⊂ *U* and *V*_*d*_ ⊂ *V* that form dense subgraphs in *B*[*U*_*c*_, *V*_*d*_], *G*[*U*_*c*_], and *H*[*V*_*d*_] simultaneously, thereby capturing genuine cross-omics pathway-level interactions rather than isolated feature-level noise.

### Estimation

Because the true cross-omics association matrix *A* is not directly observed, we instead work with a surrogate matrix *W* = {*w*_*uv*_}_*u*∈*U, v*∈*V*_ derived from pairwise association evidence, for example *w*_*uv*_ = −log(*p*_*uv*_). The within-omics structures are represented analogously by weighted adjacency matrices *W* ^(1)^ on *U* and *W* ^(2)^ on *V*. Estimation then amounts to identifying subsets *U*_*c*_ ⊂ *U* and *V*_*d*_ ⊂ *V* that are simultaneously dense in the bipartite graph *B* = (*U, V, W*) and in the induced within-omics graphs *G* = (*U, E, W* ^(1)^) and *H* = (*V, F, W* ^(2)^).

Now we propose the following criterion for selecting subgraphs:

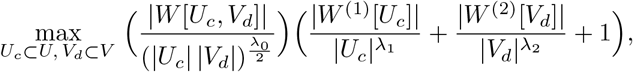

and *λ*_0_, *λ*_1_, *λ*_2_ *>* 0 control size penalties. Here *W* [*U*_*c*_, *V*_*d*_] denotes the submatrix of *W* indexed by rows *U*_*c*_ ⊂ *U* and columns *V*_*d*_ ⊂ *V*, with 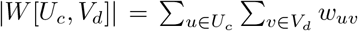 denoting the sum of all entries, where *w*_*uv*_ = − log *p*_*uv*_; |*W* ^(1)^[*U*_*c*_]| and |*W* ^(2)^[*V*_*d*_]| denote the analogous entry sums over the within-omics edge weight matrices restricted to *U*_*c*_ and *V*_*d*_ respectively. Equivalently, we write

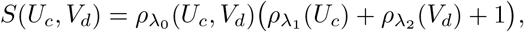

where

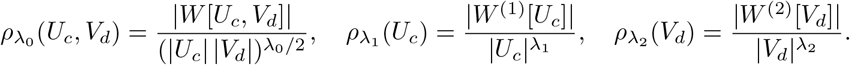

The multiplicative form gives priority to the cross-omics term while still rewarding within-omics coherence. This prevents modules driven only by strong within-layer correlation from being selected in the absence of meaningful cross-layer association.

In practice, strong within-omics correlations are common, but they do not necessarily imply meaningful cross-omics relationships. For instance, tightly co-expressed gene clusters may reflect housekeeping or other coordinated transcriptional programs without showing substantial coupling to metabolomic or proteomic variation. Likewise, groups of metabolites can be strongly correlated because they participate in the same biochemical pathway, yet still exhibit limited association with downstream gene-expression or protein-level changes. Distinguishing such within-layer coherence from biologically relevant cross-layer coupling is therefore essential, because the goal of trans-omics integration is not simply to recover internal correlation structure, but to identify cross-omics modules with interpretable biological significance.

An innovative feature of this proposed method lies in its capacity to identify one-to-multiple cluster mappings between different types of omics data. Specifically, a single cluster identified in Omics I may correspond to multiple, relatively independent clusters in Omics II. This capability allows for a more nuanced representation of biological relationships compared to conventional methods, which typically aggregate all Omics II measures into one large cluster, thereby obscuring important biological heterogeneity. For instance, in genomics-to-transcriptomics analyses, a single genomic regulatory region can influence multiple independent clusters of gene expression, each reflecting distinct functional pathways such as metabolic processes, immune responses, or developmental signaling. Similarly, within proteomics- to-metabolomics contexts, a single cluster of metabolic enzymes may independently regulate several distinct metabolite groups, including lipid, carbohydrate, and amino acid metabolism, rather than influencing a unified metabolite cluster. Thus, by capturing these biologically meaningful one-to-multiple mappings, our method provides a more accurate and comprehensive representation of complex trans-omics interactions.

#### Remark 1

(Role of the regularization parameters) For the cross-omics bipartite subgraph, the raw weight is 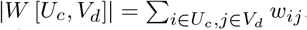. If we just maximize this sum, one will trivially get the entire matrix (*U*_*c*_ = *U, V*_*d*_ = *V*), large sets have large totals. To prevent that, we normalize by some function of |*U*_*c*_| and |*V*_*d*_|, so that we are looking for “dense” blocks, not just “large” blocks. The normalized quantity used is:

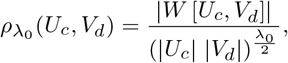

where the denominator involves |*U*_*c*_| |*V*_*d*_|, which is the number of possible edges in the bipartite subgraph.

Common choices of exponent *λ*_0_*/*2 includes: (1) *λ*_0_ = 2, then the denominator is exactly |*U*_*c*_| |*V*_*d*_|, i.e., you are normalizing by the total number of edges, turning the numerator into an average weight per edge; (2) *λ*_0_ *<* 2, the penalty is weaker: larger blocks are not fully normalized, so you bias toward finding larger but still relatively dense subgraphs; (3) *λ*_0_ *>* 2, the penalty is stronger: you prefer smaller but very dense subgraphs. The factor of 1*/*2 keeps the tuning parameter *λ*_0_ on the same scale as *λ*_1_, *λ*_2_. To see this, for the within-omics parts:

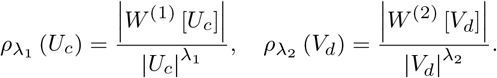

Here the natural denominator is |*U*_*c*_|^2^ or |*V*_*d*_|^2^ (since unipartite graphs have 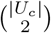 possible edges). Thus, *λ*_1_ = *λ*_2_ = 2.

#### Remark 2

(Why the additive constant is included) In the objective function:

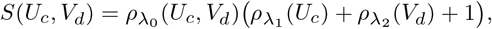

the “+1” ensures that even if intra-omics correlations are weak, the cross-omics density still contributes meaningfully to the objective. It prevents the intra-omics terms from zeroing out or overshadowing the main signal, making cross-omics detection always the baseline, with intra-omics playing a supportive boosting role.

### Algorithm

Direct optimization is combinatorial and therefore computationally challenging. We therefore use a greedy two-step strategy. Step I identifies a dense cross-omics block in the bipartite graph, which is the primary signal target. Step II then refines the retained features within each omics layer so that the final module is not only strongly associated across layers but also internally coherent within each modality.

#### Step I: Cross-omics block detection

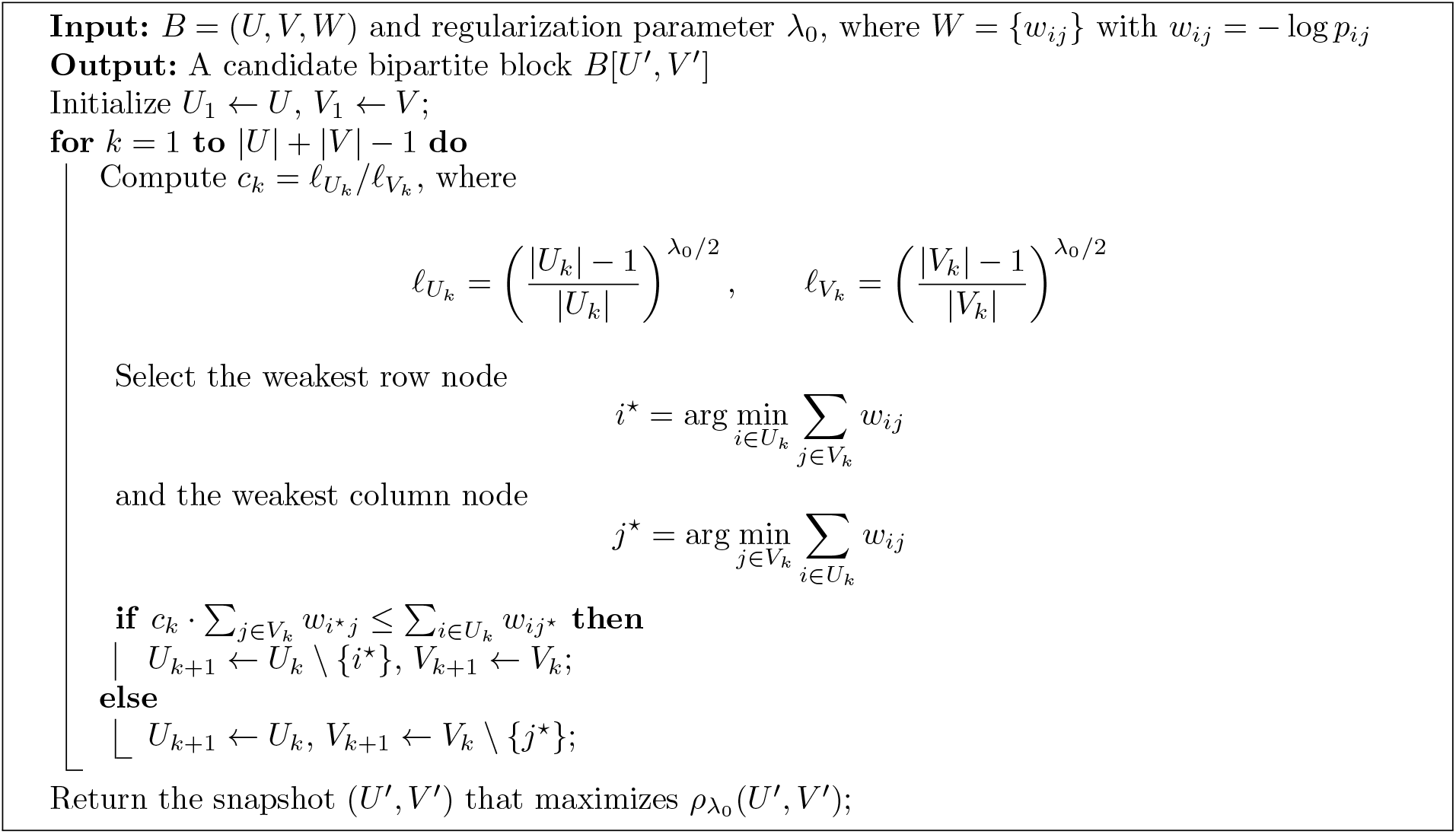

#### Step II: Within-omics refinement

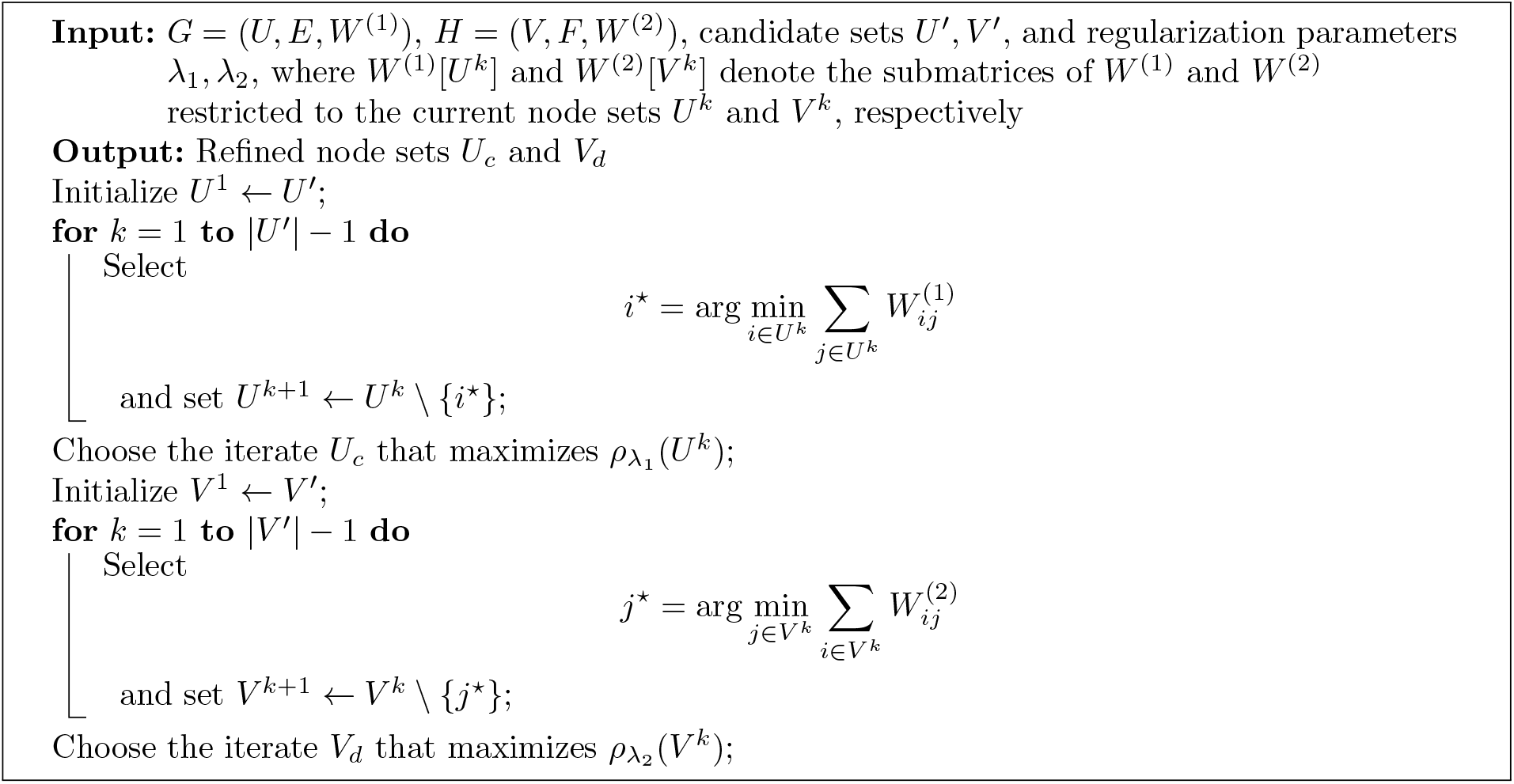

#### Algorithmic rationale

The algorithm is organized so that the primary target is identified first and then refined. Step I searches the bipartite graph for a dense cross-omics block by iteratively removing the row or column node whose deletion least harms the penalized cross-omics density and retaining the snapshot that maximizes 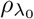. This peeling strategy favors subsets with strong collective association rather than isolated high-weight edges.

Step II starts from the feature sets retained in Step I and refines them separately within each omics layer. The aim is not to detect arbitrary within-omics clusters, but to ensure that the cross-omics block selected in Step I is also internally coherent in both modalities. The final module is therefore defined by the refined node sets *U*_*c*_ and *V*_*d*_ together with the induced cross-omics block *W* [*U*_*c*_, *V*_*d*_]. The two refinement steps are independent of each other, which naturally accommodates one-to-multiple module mappings. The Omics I peeling is run once on *U* ^*′*^ to yield a fixed *U*_*c*_; the Omics II peeling then iterates on the residual of *V* ^*′*^: after 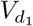 is identified and masked from *H*, the peeling is rerun on 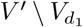 to recover 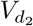, and so on until no dense block remains in the residual. Since *U*_*c*_ is held fixed throughout, this produces a collection of module pairs 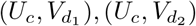 from a single candidate block (*U* ^*′*^, *V* ^*′*^), where one Omics I module is paired with multiple structurally distinct Omics II modules without imposing a one-to-one matching constraint.

A detailed step-by-step walk-through of the algorithm, including the logic of the node-removal updates and the practical interpretation of the two-stage refinement, is provided in the Supplementary Methods. The computational complexity of the cross-omics peeling step is *O*(|*U*|(|*U*| + |*V*|)). Here |*U*| = *m* and |*V*| = *n* denote the numbers of Omics I and Omics II features, respectively. Under suitable signal conditions, the probability of incorrect edge assignment decreases as sample size increases.

##### Remark 3

(Interpretation of the balancing factor *c*_*k*_) Define the denominator shrink factors caused by removing one node on each side:

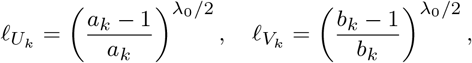

where *a*_*k*_ = |*U*_*k*_| and *b*_*k*_ = |*V*_*k*_|. These factors describe how the denominator of 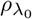 changes after removing a row or a column. The ratio

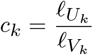

therefore rebalances the row and column degrees so that they are compared on the same penalized scale. If *U*_*k*_ and *V*_*k*_ are very different in size, removing a node from one side changes the denominator differently from removing a node from the other side; *c*_*k*_ corrects for that imbalance and makes the greedy choice consistent with the objective function.

## Supplementary Information

Supporting theoretical arguments for the dense-subgraph formulation and the greedy peeling procedure are provided in the Supplementary Methods. These supplementary derivations clarify the concentration bounds underlying recovery of dense cross-omics and within-omics subgraphs and sketch why, under suitable signal conditions, the probability of node misclassification decreases as sample size increases.

## Author information

### Contributions

J.Y. and L.H. contributed equally to this work. J.Y. and L.H. jointly developed the methodology, performed the analysis, and wrote the manuscript. S.C. conceived and supervised the project and revised the manuscript. All authors reviewed and approved the final manuscript. Please contact J.Y. for software requests.

## Declarations

### Availability of data and materials

The two real-data applications analyzed in this study used publicly available datasets. The inflammatory bowel disease analysis used data from the Inflammatory Bowel Disease Multi-omics Database (IBD-MDB) described by Lloyd-Price *et al*. [8]. The colorectal cancer analysis used the publicly available dataset reported by Yachida *et al*. [22]. Processed data objects and code required to reproduce the analyses are available from the TriGer GitHub repository at https://github.com/jy2941/TriGer. Additional information about data access is provided in the original publications cited above.

TriGer has been implemented in MATLAB, and the implementation together with the code used for all analyses in this manuscript is publicly available at https://github.com/jy2941/TriGer released under the [MIT/GPL-3.0/Apache-2.0] license.

### Ethics approval and consent to participate

Not applicable

### Consent for publication

Not applicable

## Acknowledgements

Not applicable

