## Supplemental for "Structured cross-omics interaction discovery with a triple-graph model"

### 1 Supplementary Methods

#### 1.1 Detailed algorithm walk-through

This section provides a more explicit walk-through of the two-stage optimization procedure used in the Methods section. The goal is to make clear what each step is doing, why the order of the steps matters, and how the objective function is translated into a practical algorithm.

##### 1.1.1 Overview

The method searches for a pair of feature sets, $U_c \subset U$ and $V_d \subset V$, that define a strong cross-omics association block while also remaining internally coherent within each omics layer. The optimization is carried out in two stages:

1. **Step I: cross-omics block detection.** Starting from the full bipartite graph, the algorithm iteratively removes weakly contributing nodes in order to identify a dense cross-omics block.

2. **Step II: within-omics refinement.** The feature sets retained in Step I are then refined separately in Omics I and Omics II so that the final module remains coherent in the two within-omics graphs.

This ordering is intentional. The primary signal target is the cross-omics block, whereas within-omics structure acts as a refinement criterion. If within-omics clustering were carried out first, the algorithm could prioritize highly correlated but biologically irrelevant within-layer structure that has little cross-omics relevance.

##### 1.1.2 Step I: cross-omics peeling

For any candidate bipartite block $(U', V')$, the cross-omics objective is

$$\rho_{\lambda_0}(U', V') = \frac{|W[U', V']|}{(|U'||V'|)^{\lambda_0/2}},$$

where

$$|W[U', V']| = \sum_{u \in U',\, v \in V'} w_{uv}.$$

The numerator aggregates cross-omics evidence inside the candidate block, whereas the denominator penalizes large blocks that are not sufficiently dense.

The algorithm starts from the full graph and considers removing either one node from $U_k$ or one node from $V_k$ at each iteration. For the current node sets $U_k$ and $V_k$, define

$$a_k = |U_k|, \qquad b_k = |V_k|,$$

$$r_u^{(k)} = \sum_{v \in V_k} w_{uv}, \qquad c_v^{(k)} = \sum_{u \in U_k} w_{uv},$$

where $r_u^{(k)}$ and $c_v^{(k)}$ are the current weighted row and column degrees. The weakest row and column nodes are

$$u^\star = \arg\min_{u \in U_k} r_u^{(k)}, \qquad v^\star = \arg\min_{v \in V_k} c_v^{(k)}.$$

To derive the update rule, let

$$N_k = |W[U_k, V_k]| = \sum_{u \in U_k,\, v \in V_k} w_{uv}$$

denote the total edge weight in the current bipartite block. If we remove a row node $u \in U_k$, the numerator decreases by its row degree $r_u^{(k)}$, so the resulting objective becomes

$$\rho_{\lambda_0}^{\text{row}}(u) = \frac{N_k - r_u^{(k)}}{\big((a_k - 1)b_k\big)^{\lambda_0/2}}.$$

Likewise, if we remove a column node $v \in V_k$, the updated objective is

$$\rho_{\lambda_0}^{\text{col}}(v) = \frac{N_k - c_v^{(k)}}{\big(a_k(b_k - 1)\big)^{\lambda_0/2}}.$$

The greedy step should therefore compare the best candidate row deletion and the best candidate column deletion and keep the one giving the larger next-step objective.

If a row is removed, the denominator changes differently from the case in which a column is removed. To account for this asymmetry, we define

$$\ell_{U_k} = \left(\frac{a_k - 1}{a_k}\right)^{\lambda_0/2}, \qquad \ell_{V_k} = \left(\frac{b_k - 1}{b_k}\right)^{\lambda_0/2},$$

so that

$$\begin{aligned} \big((a_k - 1)b_k\big)^{\lambda_0/2} &= (a_k b_k)^{\lambda_0/2} \ell_{U_k}, \\ \big(a_k(b_k - 1)\big)^{\lambda_0/2} &= (a_k b_k)^{\lambda_0/2} \ell_{V_k}. \end{aligned}$$

Substituting these identities gives

$$\rho_{\lambda_0}^{\text{row}}(u) = \frac{N_k - r_u^{(k)}}{(a_k b_k)^{\lambda_0/2} \ell_{U_k}}, \qquad \rho_{\lambda_0}^{\text{col}}(v) = \frac{N_k - c_v^{(k)}}{(a_k b_k)^{\lambda_0/2} \ell_{V_k}}.$$

Since the common factor $(a_k b_k)^{\lambda_0/2}$ is the same in both expressions, row deletion is preferred to column deletion if and only if

$$\frac{N_k - r_u^{(k)}}{\ell_{U_k}} \geq \frac{N_k - c_v^{(k)}}{\ell_{V_k}}.$$

Rearranging yields

$$\ell_{V_k} r_u^{(k)} \leq \ell_{U_k} c_v^{(k)}.$$

Applying this comparison to the weakest row node $u^\star$ and the weakest column node $v^\star$ gives

$$\ell_{V_k} r_{u^\star}^{(k)} \leq \ell_{U_k} c_{v^\star}^{(k)}.$$

Dividing both sides by $\ell_{V_k}$ and defining

$$c_k = \frac{\ell_{U_k}}{\ell_{V_k}}.$$

we obtain the rule used in the pseudo-code:

$$c_k \cdot r_{u^\star}^{(k)} \le c_{v^\star}^{(k)}.$$

Equivalently, since

$$r_{u^\star}^{(k)} = \sum_{j \in V_k} W_{u^\star j}, \qquad c_{v^\star}^{(k)} = \sum_{i \in U_k} W_{iv^\star},$$

the update condition can be written as

$$c_k \cdot \sum_{j \in V_k} W_{u^\star j} \le \sum_{i \in U_k} W_{iv^\star}.$$

This is exactly the criterion used in Step I: if the inequality holds, we remove the weakest row node; otherwise, we remove the weakest column node and update the graph accordingly. The role of $c_k$ is therefore not ad hoc, but to make the row-versus-column comparison consistent with the penalized objective rather than with the unnormalized edge sum.

At each iteration, the algorithm removes whichever of $u^\star$ or $v^\star$ yields the less harmful update under this objective-based comparison. The entire peeling path is recorded, and the iterate with the largest value of $\rho_{\lambda_0}(U_k, V_k)$ is retained as $(U', V')$.

##### 1.1.3 Step II: within-omics refinement

Step II refines the candidate feature sets returned by Step I. For Omics I, the within-omics objective is

$$\rho_{\lambda_1}(U_c) = \frac{|W^{(1)}[U_c]|}{|U_c|^{\lambda_1}},$$

and similarly for Omics II,

$$\rho_{\lambda_2}(V_d) = \frac{|W^{(2)}[V_d]|}{|V_d|^{\lambda_2}}.$$

The refinement step is again a peeling procedure, but it is now performed separately inside each omics layer. Starting from $U'$, the algorithm removes the node with the smallest within-omics degree in the induced graph on the current node set. The full sequence of nested subsets is recorded, and the iterate maximizing $\rho_{\lambda_1}$ is chosen as $U_c$. The same procedure is then applied to $V'$ to obtain $V_d$.

The key point is that Step II does not search the full feature space again. Instead, it refines the cross-omics candidate identified in Step I. This restriction keeps the emphasis on cross-omics signal while still enforcing within-omics coherence.

##### 1.1.4 Interpretation of the final output

The final output is a module defined by the triplet

$$(U_c,\ V_d,\ W[U_c, V_d]).$$

The first two components give the refined Omics I and Omics II node sets, and the third gives the induced cross-omics block. This representation is useful because it separates three distinct but related features of the signal: cross-layer association, within-Omics I coherence, and within-Omics II coherence.

In applications, the same framework can be used iteratively to recover multiple modules by removing or masking previously detected features and rerunning the algorithm on the remaining graph.

#### 1.2 Supporting theoretical arguments

The proofs were intended to justify two closely related claims: first, that large dense blocks are unlikely to arise from noise alone; and second, that under suitable signal conditions the peeling procedure preferentially retains nodes belonging to the true signal block.

##### Proof Part I

This part addresses the following question: *under a background-only model, how likely is it to observe a large candidate module that appears dense both across omics layers and within each omics layer?* The purpose is to justify the basic screening principle of TriGer: a module that is simultaneously dense in all three graphs should be rare under noise alone.

**Null graph setup.** Let

$$G = (U, E), \qquad H = (V, F)$$

denote the two within-omics graphs, and let

$$B^*(U, V; A)$$

denote the bipartite cross-omics graph. Under the null, suppose that the background edge probabilities are $p_1$ for $G$, $p_2$ for $H$, and $p_0$ for $B^*$. A candidate module is a pair $(U_c, V_d)$ with $U_c \subset U$ and $V_d \subset V$.

We focus on candidate blocks that are denser than background. In the bipartite layer this means

$$\frac{\sum_{i \in U_c,\, j \in V_d} h(i,j)}{|U_c||V_d|} \geq \gamma_0, \qquad \gamma_0 \in (p_0, 1),$$

where $h(i,j)$ denotes the edge indicator or weight. The condition $\gamma_0 > p_0$ means that the observed cross-omics density is stronger than expected under the null.

**Step 1: a large dense bipartite block is already unlikely.** Suppose $|U| = a$ and $|V| = b$. Let $a_0$ and $b_0$ be lower bounds on the sizes of $U_c$ and $V_d$, with

$$a_0, b_0 = \Omega(\max\{a^\epsilon, b^\epsilon\}), \qquad 0 < \epsilon < 1.$$

Then, by the dense-subgraph tail bound in Wu et al. (2021), for sufficiently large $a$ and $b$, if

$$\zeta(\gamma_0, p_0) a_0 \geq 8 \log b, \qquad \zeta(\gamma_0, p_0) b_0 \geq 8 \log a,$$

we have

$$\mathbb{P}\big(|U_c| \geq a_0,\ |V_d| \geq b_0\big) \leq 2ab \cdot \exp\left(-\frac{1}{4}\zeta(\gamma_0, p_0) a_0 b_0\right),$$

where

$$\zeta(x, y) = \left\{\frac{1}{(x-y)^2} + \frac{1}{3(x-y)}\right\}^{-1}.$$

The key takeaway is that the probability decays exponentially in the candidate block size $a_0 b_0$.

**Step 2: analogous bounds hold within each omics layer.** Now consider the Omics I graph $G = (U, E)$. If a subset $U_c$ satisfies

$$\frac{\sum_{i,j \in U_c} w(i,j)}{|U_c|^2} \geq \gamma_1, \qquad \gamma_1 \in (p_1, 1),$$

then a corresponding dense-subgraph bound yields

$$\mathbb{P}(|U_c| \geq n_1) \leq 2n_1 \cdot \exp\left(-\frac{1}{4}\zeta(\gamma_1, p_1)\nu_1^2\right),$$

for sufficiently large $n_1$, where $\nu_1 = \omega(\sqrt{n_1})$ denotes the relevant subgraph-size scale. The same logic applies to the Omics II graph $H = (V, F)$ with background density $p_2$ and threshold $\gamma_2 \in (p_2, 1)$.

**Step 3: the multilayer event is rarer still.** TriGer does not target a set that is dense in just one graph. It targets a candidate pair $(U_c, V_d)$ such that

$$\frac{\sum_{i,j \in U_c} w(i,j)}{|U_c|^2} \geq \gamma_1,$$
$$\frac{\sum_{k,\ell \in V_d} w(k,\ell)}{|V_d|^2} \geq \gamma_2,$$
$$\frac{\sum_{i \in U_c,\, j \in V_d} h(i,j)}{|U_c||V_d|} \geq \gamma_3,$$

with each threshold above its corresponding null density.

Therefore, a false positive module must simultaneously satisfy three rare events:

1. unusually high within-Omics I density,
2. unusually high within-Omics II density,
3. unusually high cross-omics density.

This implies a schematic upper bound of the form

$$\mathbb{P}\big(|U_c| \geq n_1,\ |V_d| \geq n_2,\ \text{all three density conditions hold}\big) \leq C\, n_1 n_2 \exp(-c\, n_1 n_2),$$

for constants $C, c > 0$ determined by the density gaps $\gamma_r - p_r$, $r = 0, 1, 2$.

**Conclusion.** The exact constants are not the main point. What matters is that under a background-only model, the probability of observing a large candidate module that is simultaneously dense across and within omics layers decreases exponentially with module size. This provides the theoretical justification for using multilayer density as the primary screening criterion.

*Remark* 1.1. The multilayer bound above should be interpreted as a conservative upper bound rather than a sharp exact probability calculation, because the three graph layers may be dependent. It is nevertheless sufficient to support the claim that large structured false positives are rare under the null.

**Proof Part II**

This part addresses a different question: *if a true signal block exists, why should the objective function and greedy peeling procedure preferentially recover it?* Whereas Proof Part I argues that large structured modules are unlikely to arise from noise alone, Proof Part II explains why the proposed criterion is aligned with the true signal when such a signal is present.

**Population separation between signal and background.** Let $(U^*, V^*)$ denote the true cross-omics signal block. For any candidate block $(U_c, V_d)$, define its average edge weight by

$$\bar{w}(U_c, V_d) = \frac{W[U_c, V_d]}{|U_c||V_d|}.$$

Assume that the true block has higher mean density than competing non-signal blocks:

$$\mathbb{E}[\bar{w}(U^*, V^*)] = \mu_1, \qquad \mathbb{E}[\bar{w}(U_c, V_d)] \le \mu_0,$$

for candidate blocks $(U_c, V_d)$ with limited overlap with $(U^*, V^*)$, where $\mu_1 > \mu_0$. This is the basic identifiability condition.

**Step 1: empirical densities concentrate around their means.** By a Bernstein-type concentration inequality,

$$\bar{w}(U_c, V_d) = \mathbb{E}[\bar{w}(U_c, V_d)] + O_p\left(\frac{1}{\sqrt{|U_c||V_d|}}\right).$$

Equivalently, for sufficiently large candidate blocks,

$$\mathbb{P}\Big(\big|\bar{w}(U_c, V_d) - \mathbb{E}[\bar{w}(U_c, V_d)]\big| \ge t\Big) \le \exp(-C|U_c||V_d|t^2),$$

for some constant $C > 0$. Thus the empirical average density becomes increasingly stable as the block size grows.

**Step 2: signal and non-signal blocks can be separated.** Choose any threshold $\gamma$ satisfying

$$\mu_0 < \gamma < \mu_1.$$

Then concentration implies that

1. a non-signal block is unlikely to have empirical density above $\gamma$;
2. the true block is unlikely to have empirical density below $\gamma$.

Hence sufficiently large signal blocks and background blocks are separable with high probability on the basis of empirical average density.

**Step 3: why the penalized objective is needed.** The method uses

$$\rho_{\lambda_0}(U_c, V_d) = \bar{w}(U_c, V_d) \cdot (|U_c||V_d|)^{1-\lambda_0/2} = \frac{W[U_c, V_d]}{(|U_c||V_d|)^{\lambda_0/2}}.$$

This form balances block density against block size:

- if $\lambda_0 < 2$, larger blocks are penalized less heavily and are therefore favored;
- if $\lambda_0 = 2$, the criterion reduces to average density;
- if $\lambda_0 > 2$, smaller but very dense blocks are favored.

Thus $\lambda_0$ governs the tradeoff between selecting a large diffuse block and selecting a small unstable block. Under a planted-block model, there exists a useful range of $\lambda_0$ for which the true block has larger expected criterion value than competing misspecified blocks:

$$\mathbb{E}[\rho_{\lambda_0}(U^*, V^*)] > \mathbb{E}[\rho_{\lambda_0}(U_c, V_d)].$$

**Step 4: why the peeling direction is sensible.** At each iteration, the greedy peeling algorithm removes the node with the smallest marginal contribution to the current objective. Under the planted-block picture, nodes outside $(U^*, V^*)$ have smaller expected contribution than nodes inside the signal block. Therefore, background nodes are more likely to be removed early, whereas signal nodes are more likely to remain until later iterations.

This is not a full exact-recovery proof in all regimes, but it does explain why the peeling path is statistically aligned with the target structure: repeated removal of the weakest contributors should progressively enrich the candidate block for signal nodes.

**Implication for node misclassification.** Combining population separation, concentration, and objective alignment suggests that the probability of substantial node misclassification decreases rapidly. In schematic form,

$$\mathbb{P}(e_U + e_V > \alpha) \leq \exp(-cn^{\epsilon}),$$

for constants $c > 0$ and $\epsilon \in (0, 1)$, where $e_U$ and $e_V$ denote the numbers of misclassified nodes in the two layers and $n = min(n_1, n_2)$. Equivalently,

$$\mathbb{P}(e_U + e_V \leq \alpha) \to 1.$$

**Conclusion.** Proof Part I explains why large structured modules are unlikely under noise. Proof Part II complements that result by showing why, when a true signal block exists, the size-penalized objective and greedy peeling rule are aligned with recovering it.

#### 1.3 Simulation

We generate Omics I data $\boldsymbol{X}^{(d)}_{m\times 1} = (x^{(d)}_1, \ldots, x^{(d)}_m)^T$ and Omics II data $\boldsymbol{Y}^{(d)}_{n\times 1} = (y^{(d)}_1, \ldots, y^{(d)}_n)^T$ based on the following multivariate Gaussian distribution:

$$\begin{pmatrix} X^{(d)} \\ Y^{(d)} \end{pmatrix} \sim N\left[\begin{pmatrix} \mu_X \\ \mu_Y \end{pmatrix}, \begin{pmatrix} \Sigma_{X,X} & \Sigma_{X,Y} \\ \Sigma_{Y,X} & \Sigma_{Y,Y} \end{pmatrix}\right]$$

where $\begin{pmatrix} \mu_X \\ \mu_Y \end{pmatrix}$ is the mean vector of Omics I and Omics II data respectively, and $\Sigma = \begin{pmatrix} \Sigma_{X,X} & \Sigma_{X,Y} \\ \Sigma_{Y,X} & \Sigma_{Y,Y} \end{pmatrix}$ is the partitioned variance-covariance matrix. (Such a setting can be readily extended to a latent factor model, where $X = L_1U + E_m, Y = L_2U + E_n$)

For simplicity, we set $\begin{pmatrix} \mu_X \\ \mu_Y \end{pmatrix}$ as a zero vector, representing normalized data.

#### 1.4 Extension to count-valued omics data

TriGer as presented assumes continuous measurements and uses Pearson correlation to construct the within-omics graphs $G$ and $H$ and the cross-omics bipartite graph $B$. Many omics modalities including microbiome feature tables (16S or metagenomic counts), single-cell RNA-seq, and metabolomics data processed without variance-stabilizing transformation are instead count-valued, for which Pearson correlation is not the natural dependence measure. TriGer extends to count data without modification to the graph-based optimization by replacing the association-estimation step with a *correlation copula* model. For a pair of count-valued features $(X_u, Y_v)$, let $F_u$ and $G_v$ denote their marginal cdfs. The Gaussian copula posits the existence of latent continuous variables

$$\tilde{X}_u = \Phi^{-1}(F_u(X_u)), \qquad \tilde{Y}_v = \Phi^{-1}(G_v(Y_v)),$$

where $\Phi^{-1}$ is the standard normal quantile function, such that $(\tilde{X}_u, \tilde{Y}_v)$ is jointly normal with correlation $r_{uv}$. The parameter $r_{uv}$ is the *latent correlation* and is estimated by the polyserial or polychoric method depending on whether one or both variables are discrete. Concretely, given observed counts, $F_u$ is estimated empirically and $r_{uv}$ is obtained by maximizing the bivariate Gaussian log-likelihood on the probability-integral-transformed data, or equivalently via the consistent moment estimator. The latent correlation matrix $\hat{R}_{XY} = \{\hat{r}_{uv}\}$ serves as a drop-in replacement for the Pearson cross-correlation matrix $A$ in the Methods section of the main text, and within-omics latent correlations replace the Pearson within-layer correlations used to construct $G$ and $H$. Pairwise $p$-values for the cross-omics graph $B$ are obtained by testing $H_0^{uv} : r_{uv} = 0$ using the asymptotic normality of the Fisher-$z$-transformed estimator. All downstream steps, including the size-penalized density objective, the two-stage greedy algorithm, and the permutation stopping rule, are unchanged, since they operate solely on the edge-weight matrices $W$, $W^{(1)}$, $W^{(2)}$ and not on the raw data.

**Proposition** (Validity for count data)**.** Suppose $(X_u, Y_v)$ follows a Gaussian copula model with latent correlation $r_{uv}$, and that $F_u$, $G_v$ are estimated consistently. Then the latent-correlation estimator $\hat{r}_{uv}$ is consistent and asymptotically normal, the corresponding $p$-value $p_{uv}$ is asymptotically uniform under $H_0^{uv} : r_{uv} = 0$, and the weight $w_{uv} = -\log p_{uv}$ satisfies the stochastic dominance condition stated in the Standing Assumption of Proof Part I.

*Proof sketch:* Consistency and asymptotic normality of $\hat{r}_{uv}$ follow from standard semiparametric theory for Gaussian copula models under mild moment conditions on the marginals. Asymptotic uniformity of $p_{uv}$ under the null is then a direct consequence. The stochastic dominance condition holds because $r_{uv} \neq 0$ shifts the distribution of $\hat{r}_{uv}$ away from zero, increasing $-\log p_{uv}$ in expectation; under the null $r_{uv} = 0$, asymptotic uniformity of $p_{uv}$ gives $w_{uv}$ an asymptotically standard exponential distribution, matching the null described in the Standing Assumption. Since Proof Parts I and II depend only on this dominance and not on the specific distributional form of $W_{uv}$, both results carry over to the copula-estimated weights unchanged.
